# Sex-biasing influence of autism-associated *Ube3a* gene overdosage at connectomic, behavioral and transcriptomic levels

**DOI:** 10.1101/2022.10.25.513747

**Authors:** Caterina Montani, Marco Pagani, Elizabeth De Guzman, Luigi Balasco, Filomena Grazia Alvino, Alessia de Felice, Alberto Galbusera, Thomas K. Nickl-Jockschat, Pierre Lau, Noemi Borsotti, Lorenzo Mattioni, Massimo Pasqualetti, Giovanni Provenzano, Yuri Bozzi, Michael V. Lombardo, Alessandro Gozzi

## Abstract

Many neurodevelopmental conditions, including autism, affect males more than females. Genomic mechanisms enhancing risk in males may contribute to this sex-bias. The ubiquitin protein ligase E3A gene (*Ube3a*) exerts pleiotropic effects on cellular homeostasis via control of protein turnover and by acting as transcriptional coactivator with steroid hormone receptors. Overdosage of *Ube3a* via duplication or triplication of chromosomal region 15q11-13 causes 1-2% of autistic cases. Here, we test the hypothesis that increased dosage of *Ube3a* may influence autism-relevant phenotypes in a sex-biased manner. We report robust sex-biasing effects on brain connectomics and repetitive behaviors in mice with extra copies of Ube3a. These effects were associated with a profound transcriptional dysregulation of several known autism-associated genes (e.g., FMR1, SCN2A, PTEN, MEF2C, SHANK3, TSC2) as well as differentially-expressed genes identified in human 15q duplication and in autistic patients. Notably, increased Ube3a dosage also affects multiple sex-relevant mechanisms, including genes on the X chromosome, genes influenced by sex steroid hormones, downstream targets of the androgen and estrogen receptors, or genes that are sex-differentially regulated by transcription factors. These results suggest that *Ube3a* overdosage can critically contribute to sex-bias in neurodevelopmental conditions via influence on sex-differential mechanisms.

## Introduction

Early onset neurodevelopmental conditions tend to show a sex-bias, with males being more affected than females (Rutter et al., 2003). This imbalance is especially evident in the case of autism, where the male:female ratio is around 3:1 (Loomes et al., 2017). Several ideas have been proposed to explain this phenomenon (Lai et al., 2015; Werling, 2016). Multiple risk factors that enhance risk in males have been identified, including the influence of steroid hormones, (Katsigianni et al., 2019; Baron-Cohen et al., 2020) and their prenatal programming effect on sex differences in structural and functional brain development of relevance to autism (Lombardo et al., 2012, 2020; Lai et al., 2017). In contrast, evidence for a possible genetic female protective effect has been reported. Autistic females tend to show an increased burden of rare de novo variants (Sanders et al., 2012; Jacquemont et al., 2014; Satterstrom et al., 2020) as well as higher polygenic risk from inherited common variants (Antaki et al., 2021; Wigdor et al., 2021). Rare deleterious variants are also transmitted maternally at higher rates (Desachy et al., 2015; Krumm et al., 2015). Both female protective and male risk factors have been theorized to work concurrently within a multiple liability threshold model of sex-differential risk for autism (Werling and Geschwind, 2013). However, despite these theoretical underpinnings, the exact mechanisms and genetic determinants that explain sex-bias in autism are still largely unknown.

Sex-specific genetic, transcriptomic, and regulatory architectures are implicated in most diseases and complex traits (Ober et al., 2008; Bernabeu et al., 2021; Posserud et al., 2021). Together with sex-differential hormonal environments affecting mid-gestational periods (Baron-Cohen et al., 2015; Lombardo et al., 2020), genetic risk factors may interact with sex to produce differential multi-omic effects (e.g., at transcriptome, connectome, phenome levels) that could either amplify risk in males or reduce risk in females, and thereby result in a sex-bias in autism. A key mechanism that may exert such sex-differential multi-omic effects in autism may reside within the function of the ubiquitin protein ligase E3A (Ube3a). *Ube3a* is located on chromosome 15q11-13, and deletions of this chromosomal region results in Prader-Willi or Angelman syndrome, depending on whether the paternal or maternal copy is deleted (Takumi and Tamada, 2018). Duplication or triplication of this region also has an important neurodevelopmental impact, resulting in intellectual disability, epilepsy, and autism – a syndrome commonly referred to as dup15q syndrome (Finucane et al., 1993; Urraca et al., 2013; Frohlich et al., 2019). These genetic alterations can explain 1-2% of all autism cases, and thus represent one of the strongest genetic risk factors for autism (Abrahams and Geschwind, 2008; Hogart et al., 2010). In keeping with this, animal studies have shown the increased *Ube3a* dosage reconstitutes autism-like traits in animals, an effect that may be mediated by impaired glutamatergic transmission (Smith et al., 2011).

Ube3a is commonly known for its role in protein degradation, and numerous proteins involved in neurodevelopment and autism have been reported to be a ubiquitination target of this protein, including TSC2 (Sun et al., 2015a), Ephexin5 (Margolis et al., 2010), SK2 (Sun et al., 2015b) and XIAP (Khatri et al., 2018). However, a less investigated independent function (Nawaz et al., 1999; Li et al., 2006) through which Ube3a can affect brain development is its role as transcriptional co-activator with steroid hormone receptors (El Hokayem and Nawaz, 2014). Steroid hormone receptors are known to affect developmental mechanisms related to autism (Hu et al., 2011; Sarachana and Hu, 2013; Quartier et al., 2018; Lombardo et al., 2020; Willsey et al., 2021; Gegenhuber et al., 2022; Kelava et al., 2022) and thereby represent one possible mechanistic avenue for explaining sex-bias in neurodevelopmental disorders. Through these functions, Ube3a can thus affect the transcriptomic and proteomic architecture of the developing brain (Kühnle et al., 2013; LaSalle et al., 2015; Vatsa and Jana, 2018) and may serve as a putative effector of sex-specific phenotypes of relevance to autism. Previous work supports the mechanistic plausibility of this framework, as gene expression analysis in predominantly male samples has shown convergence of cortical transcriptome dysregulation in idiopathic autism and dup15q syndrome (Parikshak et al., 2016).

Here we test the hypothesis that increase dosage of Ube3a may exert a sex-biasing influence on autism-relevant phenotypes. We used the Ube3a2X mouse model (Smith et al., 2011; Krishnan et al., 2017; Khatri et al., 2018), mimicking maternally inherited 15q11-13 triplication, to investigate how such a genomic risk factor interacts with sex to produce differential effects at connectomic, phenomic, and transcriptomic levels. Ube3a2X mice harbor two extra-copies of Ube3a transgene and exhibit deficits in cortical excitatory transmission, together with core autism traits of relevance for dup15q syndrome (Smith et al., 2011; Krishnan et al., 2017). We found that Ube3a can critically contribute to multi-omics sex-bias via transcriptional influence on genes located on the X chromosome and downstream targets of the androgen receptor, including multiple high confidence autism-associated genes. Our results uncover a powerful sex-biasing genomic influence of Ube3a that could explain some of the sex-bias in autism and related neurodevelopmental disorders.

## Results

### *Ube3a* gene dosage affects prefrontal and hypothalamic functional connectivity in a sex-specific dependent fashion

Robust alterations in brain anatomy and functional connectivity have been described in multiple autism mouse models, including mouse lines harboring genetic alterations associated with dup15q syndrome (Ellegood et al., 2014; Copping et al., 2017; Bertero et al., 2018; Pagani et al., 2021). The observed anatomical and functional alterations partly recapitulate abnormalities observed in patient populations and are thus considered a sensitive marker of developmental dysfunction. To investigate whether *Ube3a* dosage exerts sex-specific effects on brain circuits, we first carried out spatially unbiased rsfMRI connectivity mapping in male and female mice with increased *Ube3a* gene dosage (Ube3a2X), modelling dup15q syndrome (Smith et al., 2011).

Using weighted degree centrality as a metric of global connectivity (Bertero et al., 2018), we identified foci of global hypo-connectivity in the hypothalamus and thalamus of Ube3a2X mice, irrespective of sex (*t* > |2.1|, cluster-corrected, Figure 1a). Interestingly, sex-specific effects were also found with significant sex*genotype interactions in the basal forebrain and hypothalamic regions (*t* > |2.1|, cluster-corrected, Figure 1b). As shown in Figure 1b, this interaction effect reflected reduced global connectivity in Ube3a2X females (p = 0.03, Figure 1b). In contrast, Ube3a2X males displayed a non-significant trend for increased functional connectivity in both prefrontal and hypothalamic areas (p = 0.10, Figure 1b).

**Figure 1.**
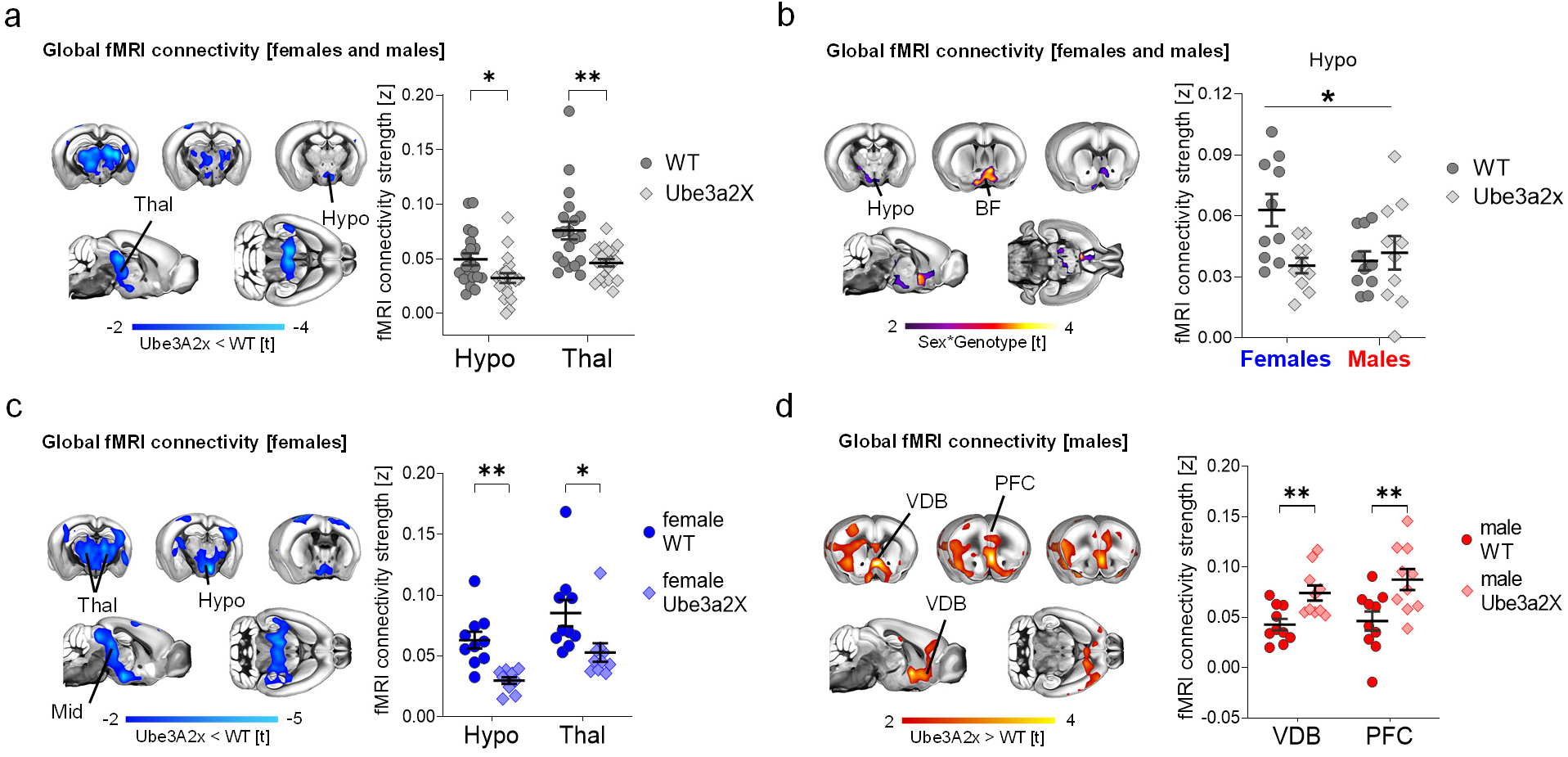
Increased Ube3a dosage affects global fMRI connectivity in a sex-dependent manner. **a)** Contrast maps (left panel) illustrating difference in global fMRI connectivity strength between WT (n = 20) and Ube3a2X (n = 20) animals, irrespective of sex (blue indicates reduce connectivity, t-test, t > 2; FWE cluster-corrected). Panel on the right illustrates quantification of global fMRI connectivity strength in representative regions of interest (t-test; Hypothalamus, t = 2.44, p = 0.019; Thalamus, t = 3.36, p = 0.002). **b)** Contrast maps (left panel) showing areas exhibiting sex*genotype interaction in global fMRI connectivity strength (purple and yellow indicates areas with significant interaction, t > 2; FWE cluster-corrected). Panel on the right illustrates the quantification of sex*genotype interaction in the hypothalamus (ANOVA’s interaction, F = 5.85, p = 0.02). **c)** Contrast maps (left panel) showing areas exhibiting decreased global fMRI connectivity strength in female Ube3a2X mice (n=10) compared to female WT (n = 10) littermates (t-test, t > 2; FWE cluster-corrected). The plot on the right illustrates quantification of global fMRI connectivity strength in representative regions of interest (t-test; Hypothalamus, t = 4.50, p < 0.001; Thalamus, t = 2.44, p = 0.026). **d)** Contrast maps (left panel) showing regions exhibiting increased global fMRI connectivity strength in male Ube3a2X mice (n=10) compared to male WT (n = 10) littermates (red indicates increased connectivity, t-test, t > 2; FWE cluster-corrected). Panel on the right illustrates quantification of global fMRI connectivity strength in both groups of males in representative regions of interest (t-test; VDB, t = 3.33, p = 0.004; PFC, t = 2.90, p = 0.009). BF, Basal Forebrain; Hypo, Hypothalamus; PFC, Prefrontal Cortex; Thal, Thalamus; SS, somatosensory cortex; VDB, Ventral Diagonal Band. *p < 0.05, **p < 0.01, FWE: family-wise error. Error bars indicate SEM.

Given the presence of sex*genotype interactions, we next carried out follow-up analyses in male and female mice, separately. These analyses revealed reduced global connectivity across a large set of mid-brain, hypothalamic, thalamic and sensory areas in Ube3a2x females (*t* > |2.1|, cluster-corrected, Figure 1c). In contrast, Ube3a2X males showed increased global connectivity in prefrontal and basal forebrain areas (*t* > |2.1|, cluster corrected, Figure 1d). To probe the circuitlevel substrates differentially affected in the two sexes, we next performed a set of seed-based connectivity analyses in regions exhibiting global connectivity differences (Figure 2). These investigations revealed foci with significant sex*genotype interactions in hypothalamic, basal forebrain and medial prefrontal areas, corroborating the involvement of these areas as key substrates for sex-divergent functional dysconnectivity produced by increased *Ube3a dosage (t* > |2.1|, FWER cluster-corrected, Figure 2a-b). Further investigation of these effects in each sex separately (Figure 2c-f) revealed that in Ube3a2X females, hypothalamic areas exhibit prominent hypo-connectivity with somatosensory cortex, thalamus and hippocampus (*t* > |2.1|, FWER cluster-corrected, Figure 2c,e). In contrast, Ube3a2X males were characterized by functional hyper-connectivity between the medial prefrontal cortex, the anterior insula, thalamic and hypothalamic regions (*t* > |2.1|, FWER cluster-corrected, Figure 2d,f). These findings suggest that hypothalamic and prefrontal circuits exhibit sex-specific, divergent patterns of dysconnectivity in mice with increased dosage of *Ube3a*.

**Figure 2.**
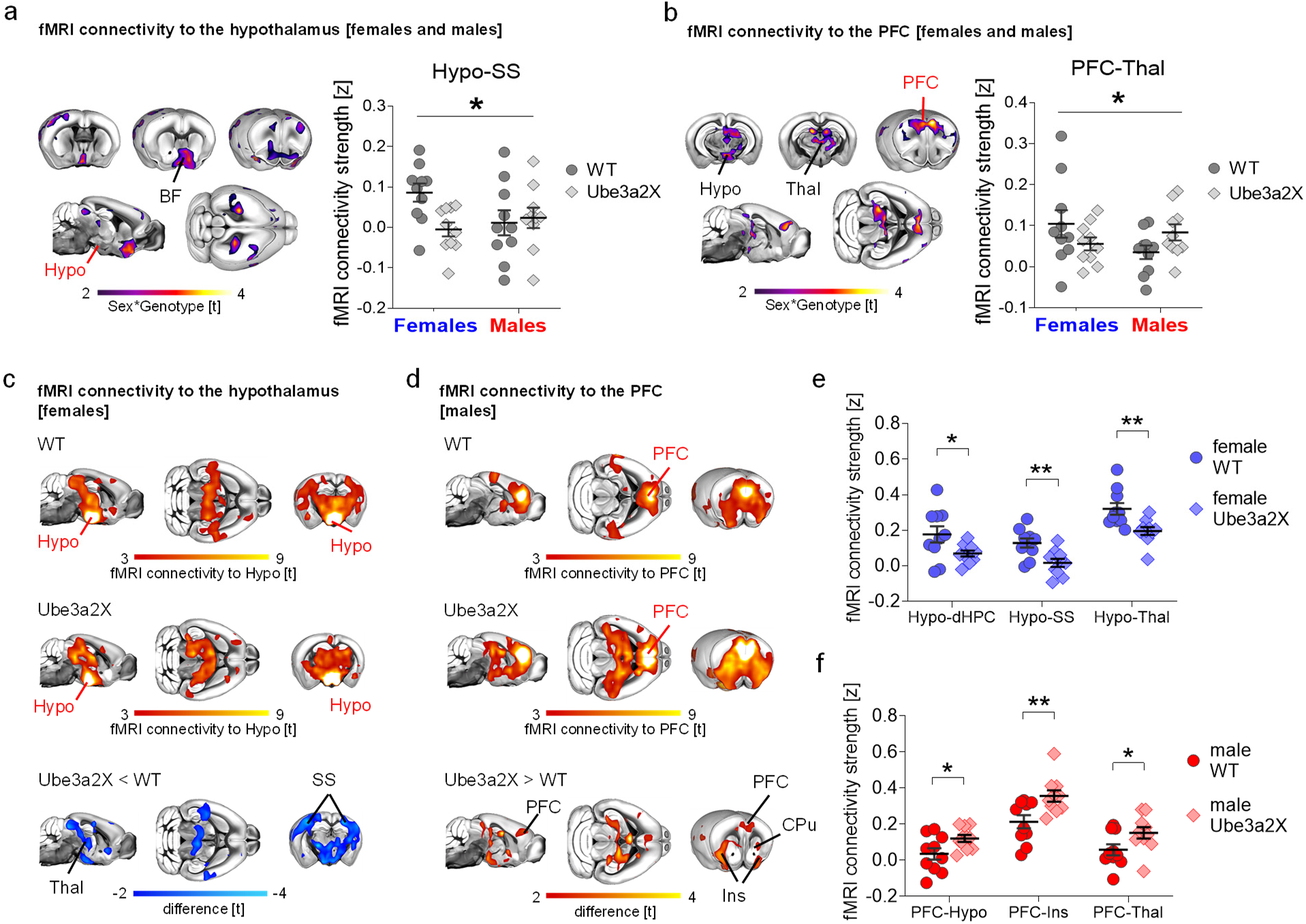
Divergent fMRI connectivity profiles in male and female Ube3a2X mutants. **a)** Seed-based fMRI connectivity mapping of the hypothalamus in Ube3a2X mice (Ube3a2X, n = 20 vs. WT n = 20, n = 10 males and females within each group). Contrast maps show areas exhibiting sex*genotype interaction of fMRI connectivity to the hypothalamus (purple and yellow coloring, GLM’s interaction, t > 2; FWE cluster-corrected). The plot (right panel) illustrates the quantification of sex*genotype interaction of fMRI connectivity strength between the hypothalamus and the somatosensory cortex (Hypo-SS, ANOVA’s interaction, F = 4.42, p = 0.04). **b)** Seed-based fMRI connectivity mapping of the prefrontal cortex in Ube3a2X mice (Ube3a2X, n =20 vs. WT n = 20, n = 10 males and females within each group). Contrast maps show areas exhibiting sex*genotype interaction of fMRI connectivity to the prefrontal cortex (purple and yellow coloring, GLM’s interaction, t > 2; FWE cluster-corrected). The plot (right panel) illustrates the quantification of sex*genotype interaction of fMRI connectivity strength between the prefrontal cortex and the thalamus (PFC-Thal, ANOVA’s interaction, F = 4.62, p = 0.04). **c)** Seed-based fMRI connectivity mapping of the hypothalamus in female WT (n = 10) and female Ube3a2X mice (n = 10). Red-yellow coloring represents regions exhibiting fMRI connectivity with the hypothalamic seed region (red lettering) in female control (WT, top panel, one sample t-test, t > 3) and female Ube3a2X (Ube3a2X, middle panel, one sample t-test, t > 3) mice. The corresponding contrast map (Ube3a2x < WT) is reported at the bottom of the panel (blue indicates reduced connectivity, t-test, t > 2). All statistics are FWE cluster-corrected. **d)** Seed-based fMRI connectivity mapping of the prefrontal cortex in male WT (n = 10) and sex-matched Ube3a2X mutants (n = 10). Red-yellow coloring represents regions exhibiting rsfMRI functional connectivity with the prefrontal seed region (red lettering) in male control (WT, top panel, one sample t-test, t > 3) and male Ube3a2X (Ube3a2X, middle panel, one sample t-test, t > 3) mice. The corresponding contrast map (Ube3a2x > WT) is reported at the bottom of the panel (red indicates increased connectivity, t-test, t > 2). All statistics are FWE cluster-corrected. **e)** Quantification of fMRI connectivity strength between the hypothalamus and dorsal hippocampus (Hypo-dHPC, unpaired t-test; t = 2.21, p = 0.048), somatosensory cortex (Hypo-SS, unpaired t-test; t = 3.26, p = 0.004) and thalamus (Hypo-Thal, unpaired t-test; t = 3.14, p = 0.006) in the two female groups. **f)** Quantification of fMRI connectivity strength between the anterior cingulate and dorsal hypothalamus (PFC-Hypo, unpaired t-test; t = 2.27, p = 0.038), insula (PFC-Ins, unpaired t-test; t = 2.96, p = 0.008) and thalamus (PFC-Thal, unpaired t-test; t = 2.15, p = 0.045) in the two male groups. BF, Basal Forebrain; dHPC, dorsal hippocampus; Hypo, Hypothalamus; Ins, Insula; PFC, Prefrontal cortex; SS, somatosensory cortex; Thal, thalamus. *p < 0.05, **p < 0.01, FWE, family-wise error. Error bars of the plots indicate SEM and each dot represents a mouse.

Local fMRI connectivity is also often disrupted in mouse models of autism (Liska et al., 2018; Pagani et al., 2019). We thus investigated if sex-biased changes in connectivity would also be detectable on a local scale. Local connectivity mapping in Ube3a2X mice revealed foci of robustly decreased local connectivity in hypothalamus, thalamus and hippocampus (*t* > |2.1|, cluster corrected, Figure S1a), irrespective of sex. Significant sex*genotype interactions were observed in prefrontal, hippocampal and hypothalamic regions (*t* > |2.1|, cluster corrected, Figure S1b). This effect was mainly driven by decreased local connectivity in Ube3a2X females (p = 0.04, Figure S1b). Interestingly, sex-specific effects were not apparent in brain anatomy (Figure S2). Gray matter (GM) voxel-based morphometry (Pagani et al., 2016a) revealed robust bilateral reductions in GM volume in the amygdala, thalamus, and hippocampus in Ube3a2X mice irrespective of sex (t > |2|, cluster-corrected, Figure S2. No sex*genotype interactions were identified upon voxelwise mapping (t > |2.1|, Figure S2c).

Taken together, these imaging studies show that hypothalamic, and prefrontal circuits exhibit divergent, sex-specific patterns of functional dysconnectivity in Ube3a2X mice.

### Increased stereotyped behavior in male but not female Ube3a2X mice

The observation of sex*genotype interactions in fMRI connectivity led us to investigate whether sex-specific behavioral dysfunction would be detectable in behavioral domains relevant to autism and other male-biased neurodevelopmental disorders. Motor issues (e.g., delays in achieving early motor milestones, hypotonia, clumsiness, difficulties across visuomotor, fine and gross motor skills) are a common, yet non-diagnostic, feature of many autistic individuals that increases with increased severity in core diagnostic domains (Bhat and Narayan Bhat, 2020; Bhat, 2021; Bhat et al., 2022). Previous research also indicates that 15qdup syndrome in humans is associated with motor impairments (Distefano et al., 2016; Wilson et al., 2020). We thus used the rotarod test to probe the presence of sex*genotype interactions in locomotor activity and motor coordination in Ube3a2X mice (Figure 3). We found that Ube3a2X mutants exhibited motor impairments as assessed with latency to fall score (F = 16.2; p < 0.001, genotype, 2-way ANOVA, Figure 3b), but this effect was not sex-specific (sex*genotype interaction, F = 0.29, p = 0.59, Figure 3c). Further investigations using the open field test (Figure S3a) revealed that Ube3a2X mice, irrespective of sex, showed comparable mobility (total distance travelled and frequency of rotations), time spent in the center of the field, and time spent wall rearing to control wild type littermates (p > 0.32, genotype, all tests, 2-way ANOVA, Figure S3a). These results rule-out the presence of prominent anxiety-like phenotypes or hyperactivity in these mutants, hence arguing against a confounding contribution of motor hyperactivity on the results obtained with the rotarod test.

**Figure 3.**
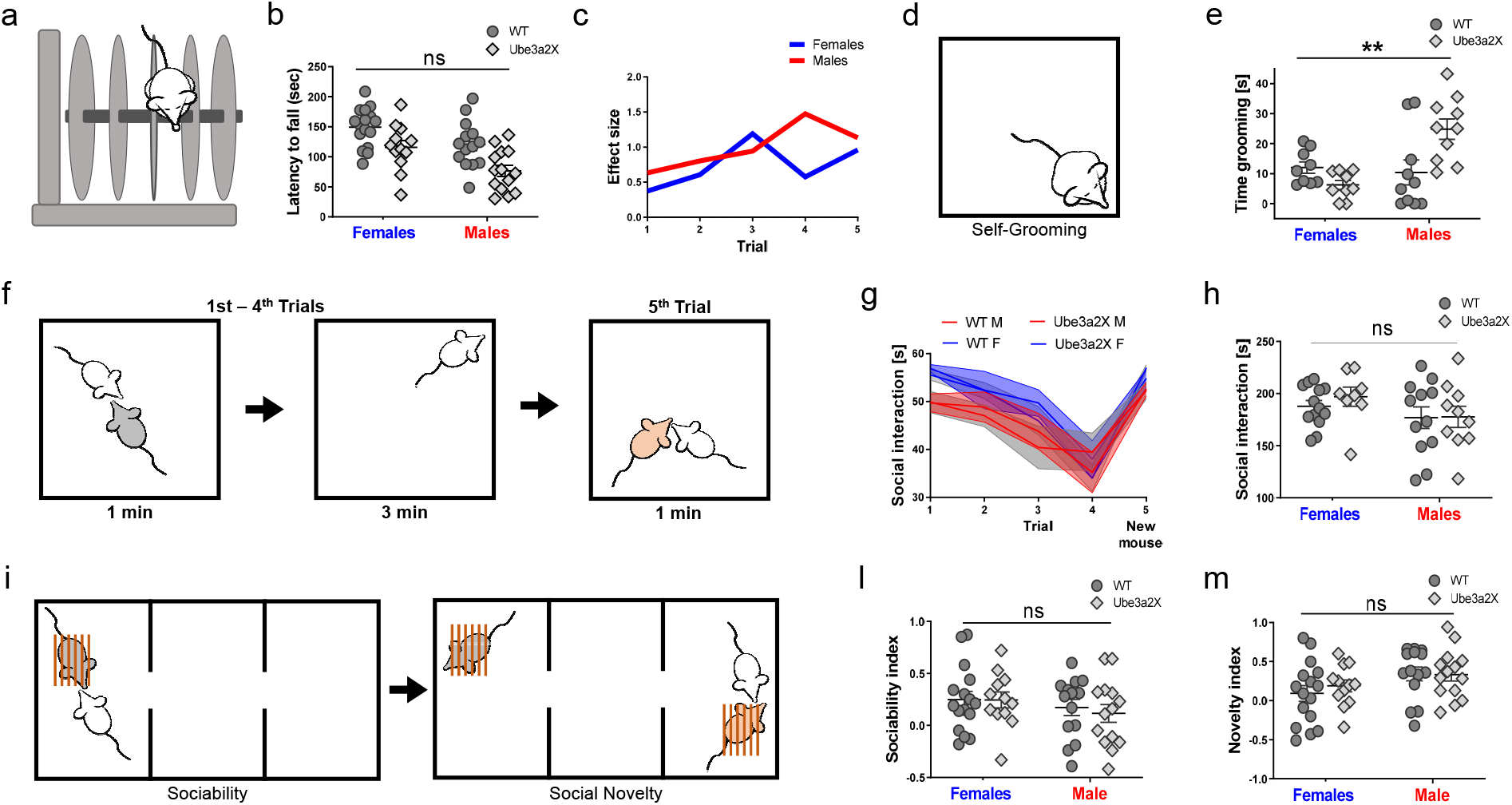
Increased Ube3a dosage affects stereotyped behavior in a sex dependent manner. **a)** Schematics of the rotarod used to assess locomotor activity. **b)**). Latency to fall was quantified for Ube3a2X mice (Ube3a2X, n = 26, n = 14 males and n = 12 females) and WT littermates (n = 30, n = 14 males and n = 16 females). Sex*genotype interaction was not significant (“ns”, ANOVA’s interaction, F = 0.29, p = 0.59). Sex and genotype factors were instead statistically significant (ANOVA, F = 12.3, p < 0.001, F = 16.2, p < 0.001, respectively) and driven by decreased latency to fall in transgenic males (post hoc test Tukey’s multiple comparisons test ***p < 0.001). **c)** Quantification of the effect size (Cohen’s d) for the latency to fall across trials in the two sexes. **d)** Schematics of the self-grooming test. **e)** Quantification of time spent grooming by Ube3a2X mice (n = 20, n = 10 males and n = 10 females) and WT littermates (n = 20, n = 10 males and females within each group). Sex*genotype interaction was statistically significant (ANOVA’s interaction F = 10.95, **p = 0.002) and driven by increased grooming in male Ube3a2x mutants. **f)** Schematics of the habituation/dishabituation social interaction test. **g)** Social interaction duration in the habituation/dishabituation test is reported for all trials (WT, n = 25, n = 12 males and n = 13 females) and Ube3a2X (n = 18, n = 10 males and n = 8 females). **h)** Cumulative social interaction duration in the first four trials of the habituation/dishabituation test. Sex*genotype interaction was not statistically significant (“ns”, ANOVA’s interaction F = 0.21, p = 0.64). **i)** Schematics of the three-chamber test. **l)** Quantification of the sociability index for WT (n = 30, n = 14 males and n = 16 females) and Ube3a2X mice (n = 26, n = 14 males and n = 12 females). Sex*genotype interaction was not significant (“ns”, ANOVA’s interaction F = 0.11, p = 0.73). **m)** Quantification of novelty index. Sex*genotype interaction was not significant (“ns”, ANOVA’s interaction F = 0.33, p = 0.57).

We next investigated the presence of autism-like stereotyped behavior using selfgrooming scoring (Silverman et al., 2010a). Interestingly, these investigations revealed robust sex*genotype interactions (F = 11.0; p = 0.002, 2-way ANOVA, Figure 3e), explained by Ube3a2X males spending more time self-grooming compared to WT male littermates (p = 0.008, Figure 3e), whereas Ube3a2X females exhibited a reverse trend of less time spent on self-grooming (p = 0.56, Figure 3e). These results show that increased stereotyped behaviors are present in male but not female mutant mice.

We next probed social behavior in control and Ube3a2X mutants using a habituation/dishabituation social interaction test (Huang et al., 2014) (Figure 3f-h). We did not observe any genotype-dependent difference in sociability in Ube3a2X mice, nor sex*genotype interactions, both in terms of interaction time with the familiar mouse, and also upon measuring interaction with a novel stimulus mouse (F = 0.78, p > 0.51, all comparisons, 2-way ANOVA, Figure 3g-h). To further investigate social behavior in Ube3a2x mice, we also tested mutant and control mice in a three-chamber test (Fig 3i-m, Figure S3b) (Silverman et al., 2010a). Also in this test, we did not find any significant genotype-dependent effects, or sex*genotype interaction in either sociability (F = 0.11; p = 0.73; 2-way ANOVA, Figure 3l) or social novelty index (F = 0.33; p = 0.57; Fig 3m). In summary, increased Ube3a dosage affects motor ability but not sociability or social habituation responses. Sex-specific effects on stereotyped behavior were apparent, indicative of autism-like increased stereotyped behaviors in male but not in female Ube3a2X mutants.

### Increased Ube3a dosage results in sex-specific PFC transcriptomic dysregulation

Given the global and local connectivity abnormalities converging on the medial prefrontal cortex (PFC) and hypothalamus (Hypo), we next investigated if increased *Ube3a* dosage results in sex-specific transcriptomic dysregulation in those regions. Bulk tissue from the PFC and Hypo was used to quantify gene expression with RNA-seq and analysis was tailored to identify differentially expressed (DE) genes for main effects of sex, genotype, as well as the sex*genotype interaction. In the PFC, 2,625 genes were detected as DE at FDR q<0.05 for the sex*genotype interaction. In contrast, no genes survived FDR correction for the interaction effect in Hypo (Figure S4; Supplementary Table S1). PFC sex*genotype interaction DE genes fell into two main classes – 1) so-called ‘DE-’genes, whereby the minus sign in the acronym is meant to describe downregulated expression in Ube3a2X males (e.g., WT > Ube3a2X) and 2) so-called ‘DE+’ genes, whereby the plus sign in the acronym describes upregulated expression in Ube3a2X males (e.g., Ube3a2X>WT). Since these are interaction effects, the effects observed in Ube3a2X males are reversed when considering females (Figure 4a). The main effect of genotype identified only *Ube3a* after FDR correction in both PFC and Hypo. For a list of genes that survived statistical thresholding of the main effect of sex, please see Figure S4 and Supplementary Table S2.

**Fig. 4.**
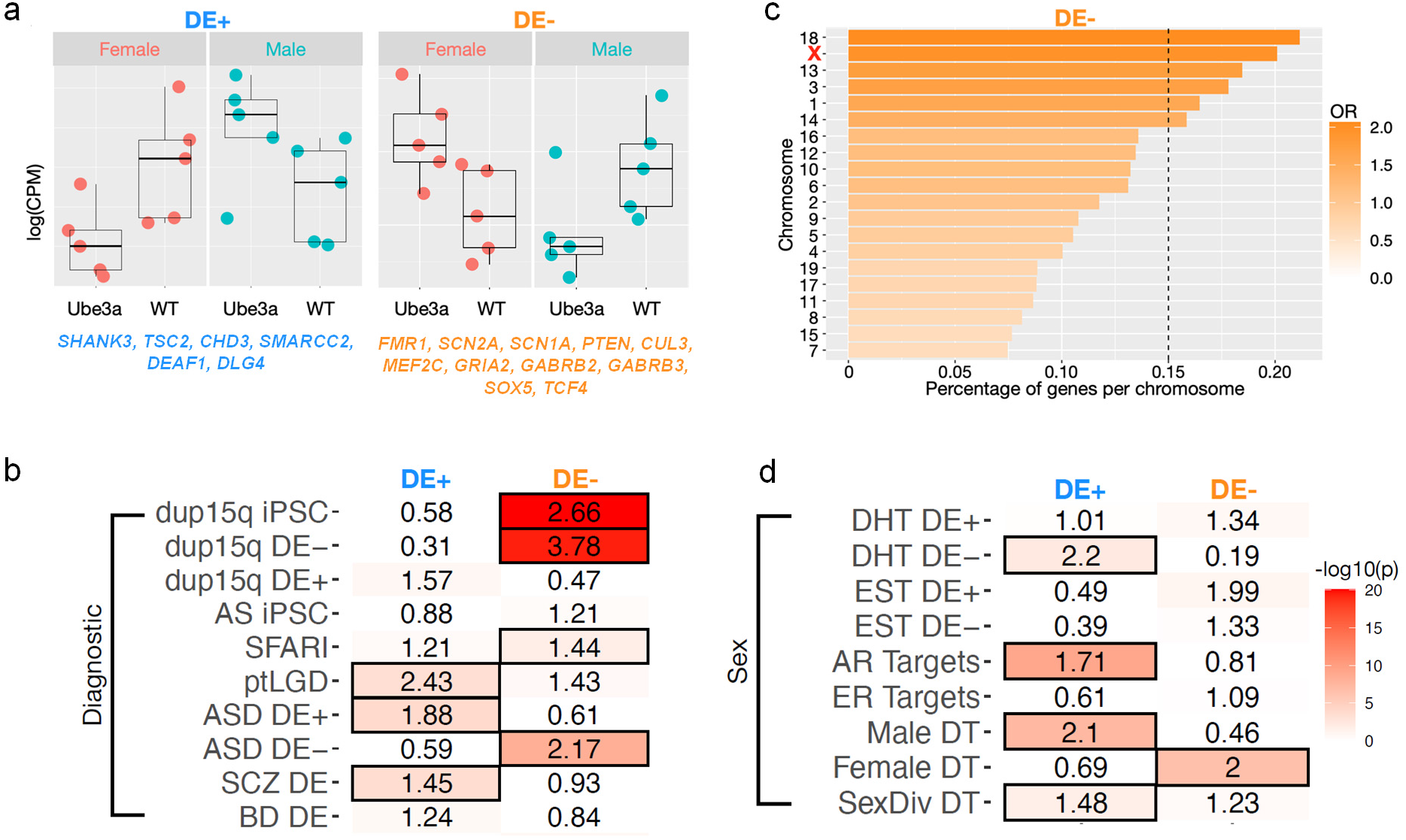
Sex-specific PFC transcriptomic dysregulation by Ube3a over-expression and enrichment with autism-associated, dup15q, and sex-relevant mechanisms. **a)** Plots show examples of DE genes for the sex*genotype interaction (DE-; light blue DE+; orange). **b)** Heatmap showing enrichments with gene lists from dup15q (dup15q DE) (from Parikshak et al., 2016), SFARI gene list, private (pt) inherited likely gene disrupting (LGD) variants (from Wilfert et al 2021), Autism Spectrum Disorders (ASD), Schizophrenia (SCZ) and Bipolar Disorder (BD) (from Gandal et al., 2018), iPSC-derived neurons from dup15q (dup15q iPSC) and Angelman Syndrome (AS iPSC) individuals (from Germain et al., 2014). The plus or minus symbols on the acronyms indicate down-(minus) or up-regulated (plus) expression. The numbers within each cell indicate the enrichment odds ratio, while color indicates the −log10(p-value) for the enrichment test. Cells outlined in black pass FDR q<0.05 for multiple comparisons correction. **c)** Plot showing the percentage of genes per each chromosome that are DE-. Color indicates the enrichment odds ratio. The X chromosome is highlighted in red. The vertical dotted line indicates the FDR q < 0.05 threshold. **d)** Heatmap showing enrichments between the sex*genotype interaction DE genes and genes relevant to sex-hormones or sex-differential gene regulation. The numbers in each cell indicate the enrichment odds ratio, and the color indicates the −log10(p-value) for the enrichment test. Cells outlined in black pass FDR q<0.05 for multiple comparisons correction. DHT DE+ and DHT DE- are genes that are up-(plus) or down-regulated (minus) in expression after dihydrotestosterone (DHT) manipulation (Lombardo et al., 2020). EST DE+ and EST DE- are genes that are up-(plus) or down-regulated (minus) in expression after treatment with estrogen (EST) (Willsey et al., 2021). AR Targets are downstream target genes of the androgen receptor (AR) as defined by ChIP-seq in human neural stem cells (Quartier et al., 2019). ER Targets are downstream target genes of the estrogen receptor (ER) (Gegenhuber et al, 2022). Male DT, Female DT, and SexDiv DT and genes that are sex-differentially targeted by transcription factors (Lopes-Ramos et al., 2020).

### Mouse-human cross-species translation via enrichment for autism-associated and dup15q genes

Having identified significant sex-specific transcriptome dysregulation in PFC, we next asked if such genes are of relevance to known genetic mechanisms of importance in human patients with either autism or dup15q syndrome. The combined set of all DE+ and DE-genes were significantly enriched for genes annotated on SFARI Gene (https://gene.sfari.org) as being associated with autism (OR = 1.57, p = 0.01). This enrichment comprised many notable high-confidence genes such as *FMR1, SHANK3, SCN2A, SCN1A, PTEN, CUL3, TSC2, MEF2C, GRIA2, GABRB2, GABRB3, CHD3, SOX5, SMARCC2, DEAF1, DLG4*, and *TCF4.* Splitting the enrichment analysis in DE+ and DE-gene sets further revealed that this SFARI enrichment was driven primarily by the DE-gene set (Figure 4b). Going beyond evidence in SFARI Gene, we also tested for enrichments with ultra-rare private inherited mutations (ptLGD) that contribute to at least 4.5% of autism risk (Wilfert et al., 2020). We found that DE+, but not DE-genes, were enriched for these ptLGD genes (Figure 4b). In line with these results, we also found that DE-genes, that are downregulated in Ube3a2X males, were enriched for genes that are downregulated in postmortem cortical tissue of a primarily male sample of human patients with autism (Gandal et al., 2018). In contrast, DE+ genes (i.e. upregulated in Ube3a2X males), were enriched for genes with upregulated expression in post-mortem cortical tissue of a predominantly male group of patients with autism (Gandal et al., 2018) (Figure 4b). DE+ genes also overlapped with dysregulated cortical transcriptome signal in patients with schizophrenia (SCZ; Figure 4b), a finding that may be expected given some overlap in the genomic mechanisms involved in autism and schizophrenia. Further underscoring the mouse-human cross-species translational value of our findings, we found that DE-genes were highly enriched for DE genes in iPSC-derived neurons from dup15q but not from Angelman syndrome individuals (Germain et al., 2014) (Figure 4b). Furthermore, DE-genes were highly enriched for genes that are downregulated in cortical tissue of human dup15q patients (Parikshak et al., 2016) (Figure 4b). For a complete statistics of each gene list used in the enrichment tests, please see Supplementary Table S3. These enrichment results support the translational relevance for human patients with autism or dup15q syndrome of the sex-specific transcriptomic dysregulation produced by increased dosage of *Ube3a.*

### Sex-specific transcriptomic dysregulation of Ube3a impacts sex-relevant genomic mechanisms

Sex differences in the brain are theorized to be mediated by mechanisms driven by genes located on the sex chromosomes - in particular on the X chromosome (Raznahan and Disteche, 2021). Thus, we next tested whether PFC sex*genotype interaction genes were disproportionately more common on specific chromosomes such as the X chromosome, than expected by chance. Indeed, we found that DE-, but not DE+ genes, were disproportionately located on several chromosomes (significant after FDR q<0.05) and that the X chromosome was one of these (Figure 4c; Figure S5).

One of the prominent roles of Ube3a is its function as transcriptional co-activator with steroid hormone receptors (e.g., *AR, ESR1, ESR2, PGR).* This suggests that Ube3a may influence transcription in a manner dependent on these sex-relevant mechanisms. To examine this hypothesis in more detail, we next investigated how PFC sex*genotype DE genes might overlap with gene lists incorporating sex-relevant mechanisms, such as genes sensitive to the transcriptional influence of sex steroid hormones, downstream targets of the androgen and estrogen receptors, or genes that are sex-differentially regulated by transcription factors. These analyses revealed that DE+ genes overlap significantly with downstream targets of the androgen receptor (AR Targets), but not targets of the estrogen receptor (ER Targets). DE+ genes were also enriched for genes downregulated by potent androgens such as dihydrotestosterone (DHT DE-).

Genes that are sex-differentially regulated by transcription factors were important as well. The DE+ set was enriched for genes that show stronger male-regulatory influence (Male DT), and for genes with relatively equal proportions of male-biased and female-biased transcription factors exerting regulatory influence (SexDiv DT). In contrast, DE-genes were significantly enriched only for genes with female-biased regulatory influence (Female DT) (Figure 4d). For a complete statistics of each gene list used in the enrichment tests, please see Supplementary Table S3. Overall, these results show that *Ube3a* over-expression impacts gene networks and systems under the influence of diverse sex-relevant mechanisms, including the effect of genes sensitive to steroid hormone influence, downstream targets of steroid hormone receptors, as well as genes that are sex-differentially targeted by transcription factors.

### Sex-specific transcriptomic dysregulation by Ube3a impacts convergent ASD-relevant biological systems and pathways

Several studies have noted common downstream biological processes/pathways and cell types that may unify the heterogeneous genomic and molecular basis behind ASD. Amongst the most important processes/pathways are synapse, transcription and chromatin remodeling, protein synthesis and translation, protein degradation, cytoskeleton processes, splicing, and numerous signaling pathways (e.g., RAS/ERK/MAPK, PIK3/AKT/ mTOR, WNT) (Bourgeron, 2015; Courchesne et al., 2019; Eyring and Geschwind, 2021). Thus, we next examined the PFC sex*genotype DE gene sets for enrichments in these processes/pathways and cell types. For biological process enrichment analysis we used GeneWalk to get context-specific and gene-level enrichments for Gene Ontology Biological Process (GO BP) terms. This analysis resulted in a variety of key ASD-relevant processes. To visualize these GO BP processes and the DE genes that go along with such enrichments within protein-protein interaction (PPI) networks, we show in Figure 5a the genes which map onto 3 clusters of terms: 1) synaptic, glutamatergic, GABAergic, ion channel proteins (green), 2) transcription and chromatin remodeling proteins, proteins within mTOR and ERK signaling pathways, as well as steroid hormone receptors (*AR*, *ESR1, ESR2, PGR*) and proteins that show enrichments for androgen and estrogen receptor signaling (blue), and 3) proteins involved in translation and protein synthesis. Amongst this PPI network, the SFARI ASD genes are noted with a black outline and the central gene manipulated here, *Ube3a*, is outlined in orange. This evidence of highly significant protein-protein interactions (actual edges = 1015, expected edges = 405, p < 1.0e-16) and highly ASD-relevant GO BP enrichment showcases a clear example of how the PFC sex*genotype DE genes are embedded within a complex systems level biological pathology that integrates abnormalities along these key processes and pathways.

### Sex-specific PFC transcriptomic dysregulation by Ube3a differentially affects neuronal and glia cell types

Finally, we asked what cell type markers are enriched within the PFC sex*genotype interaction gene set. Here we used lists of cell type markers from a mouse single cell transcriptomic atlas from the Allen Institute covering a diverse array of multiple glutamatergic and GABAergic neuronal cell types in mouse isocortex and hippocampus, as well as numerous glial and other non-neuronal cell types (Yao et al., 2021). These analyses uncovered that both DE- and DE+ gene sets were significantly enriched for a number of glutamatergic and GABAergic cell type markers. However, the magnitude and coverage of enrichments with these neuronal cell types were much stronger and broader for the DE-gene set. Setting the DE+ gene set apart from DEgenes, we also identified strong enrichments with numerous astrocyte and oligodendrocyte cell type markers, whereas no significant enrichments in these markers were present in the DE-gene set (Figure 5b). This result is suggestive of remarkable cell type specificity in how *Ube3a* overexpression drives sex-specific PFC transcriptomic dysregulation. Genes down-regulated in males, but up-regulated in females, affected diverse glutamatergic and GABAergic cell types, whereas genes up-regulated in males, but down-regulated in females affected astrocyte, oligodendrocyte, and some glutamatergic and GABAergic neuronal cell types.

**Fig. 5.**
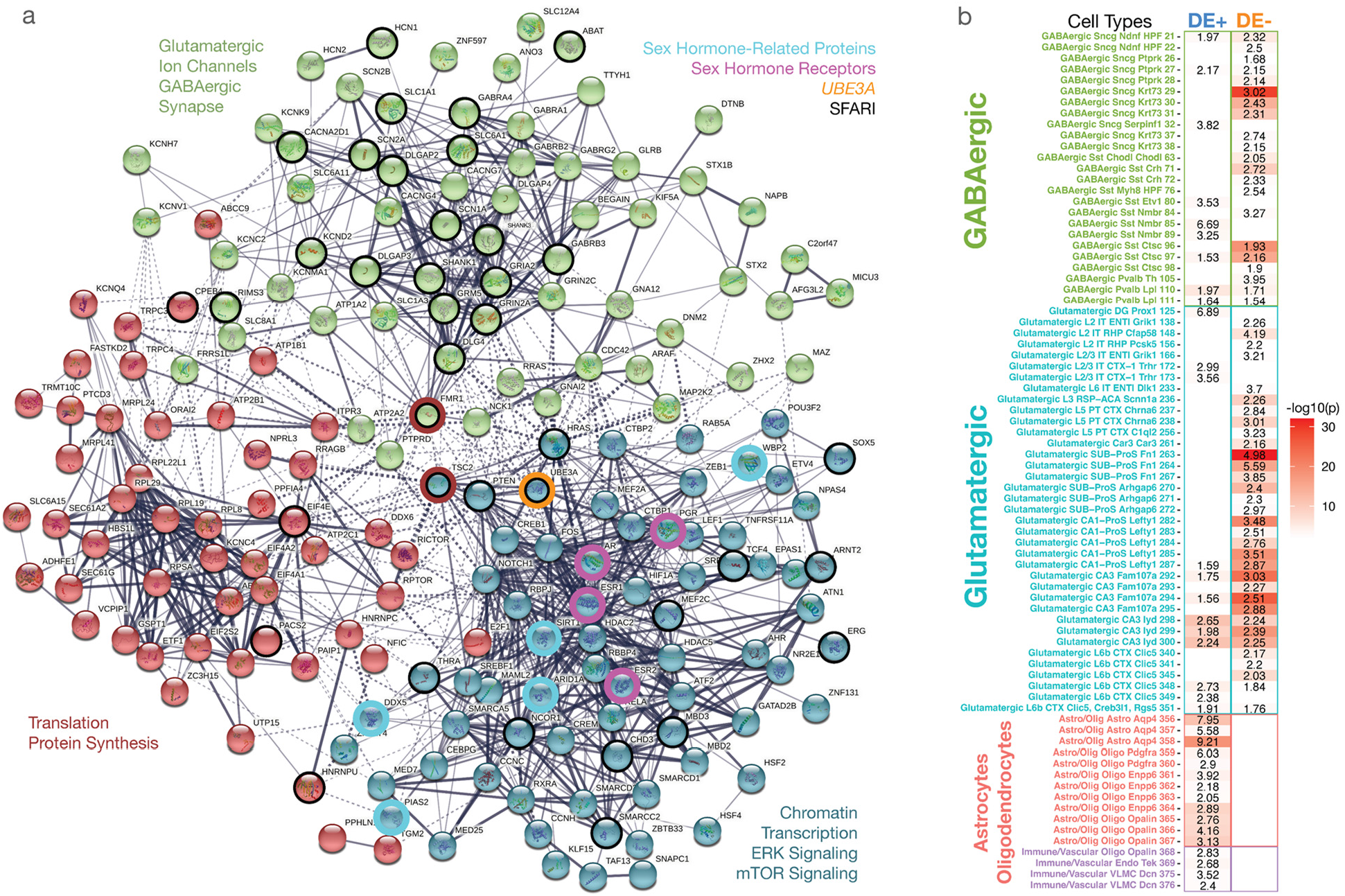
Sex-specific PFC transcriptomic dysregulation by Ube3a over-expression impacts convergent ASD-relevant biological systems, pathways, and cell types. **a)** Protein-protein interaction (PPI) graph of PFC sex-by-genotype interaction genes. Nodes are colored according to a k-means clustering solution with k=3. These clusters also segregate genes with GO BP enrichment terms specificed in text next to each cluster. Nodes are circled in black if they are SFARI ASD genes. Nodes circled in turquoise are sex-hormone related proteins, whereas magenta circled nodes are sex-hormone receptors. Ube3a is circled in orange. FMR1 and TSC2 are circled in red as they translation and protein synthesis relevant genes, but are connector hub genes in the network belonging to other clusters and which also connect the larger clusters synapse and chromatin clusters to the translation and protein synthesis cluster. **b)** Cell type enrichment heatmap showing how DE+ or DE-PFC sex-by-genotype interaction gene sets (columns) are enriched in numerous cell types markers (specified on the rows) from the Allen Institute mouse scRNA-seq data (Yao et al., 2021, Cell). DE+ genes strongly hit a variety of glutamatergic and GABAergic cell types, whereas DE-genes show specific enrichments with astrocyte and oligodendrocyte cell types. The numbers in each cell indicate the enrichment odds ratio, while the coloring indicates the −log10(p-value). Only enrichments significant at FDR q<0.05 are shown.

## Discussion

While the role of *Ube3a* mutations in determining monogenic forms of developmental disorders is well-established (Buiting, 2010), the possibility that *Ube3a* overdosage might exert sex-biasing influence has not been extensively explored. Here we document a previously unreported sex-biasing effect of *Ube3a* gene overdosing at the connectomic, behavioral and transcriptomic levels.

Our investigations revealed a sex-dependent effect of *Ube3a* overdosage on multiple (endo)phenotypes of relevance to autism, including changes in rsfMRI connectivity. Altered interareal rsfMRI coupling is a hallmark of autism, where the observed connectivity changes are prominent but also highly heterogeneous (Hull et al., 2017; Hong et al., 2020; Nair et al., 2020). Recently, fMRI-based connectivity mapping across 16 mouse mutants harboring different autism-relevant etiologies revealed a broad spectrum of connectional alteration (Zerbi et al., 2021), providing compelling evidence that heterogeneous findings in autism are likely to, at least partly, reflect the etiological heterogeneity of autism. Our results show that autism-relevant connectivity changes can also critically differ across sexes within the same etiological domain, hence underscoring a key, yet still underappreciated, dimension to the investigation of the origin and significance of functional dysconnectivity in brain disorders (Lawrence et al., 2020; Olson et al., 2020; Floris et al., 2021; Gozzi and Zerbi, 2022; Supekar et al., 2022). More broadly, these findings add to the emerging concept that autism and related neurodevelopmental disorders are characterized by a broad spectrum of connectivity alterations that are very sensitive to (and strongly biased by) the underlying etiological mechanisms (Zerbi et al., 2021). In this respect, the increased motor stereotypies and fronto-striatal connectivity in Ube3a2X we found in male mutants are of interest, as we recently described remarkably similar phenotypes in a genetically distinct mouse models (Tsc2^+/-^ (Pagani et al., 2021)). Interestingly, in the same study we also showed that analogous fronto-striatal connectivity signatures are detectable in subsets of idiopathic autism patients, where they are associated with a gene co-expression network involving mTOR-Tsc2. These findings suggest that, while etiological diversity is a prominent contributor to autism heterogeneity, congruent autism relevant circuit dysfunction may arise as a result of the dysregulation of distinct, mechanistically-dissociable gene co-expression networks.

Our work also reports a sex-invariant effect of Ube3a2X overdosage on large scale neuroanatomy. Specifically, we report reduced gray matter volume in the amygdala, thalamus and hippocampus of Ube3a2x mutants. Similar findings have been previously shown in mice overexpressing Ube3a in Camk2a positive excitatory neurons (Copping et al., 2017). Interestingly, dup15q patients have been reported to present brain morphological abnormalities, mostly involving the hippocampus (Boronat et al., 2014), hence underscoring the translational relevance of these anatomical alterations. The role of Ube3a in neuronal morphology growth and maturation has been largely investigated (Khatri and Man, 2019), but the exact relationship between Ube3a gene dosage and large scale brain anatomy remains elusive. Our findings of sex-invariant anatomical changes, but sex-specific connectivity alterations, in Ube3a2X mice suggest that Ube3a overdosage may affect brain connectivity and anatomy through distinct etiopathological cascades.

Because of the pleiotropic influence of *Ube3a* on multiple molecular and transcriptional pathways, ranging from protein degradation to transcriptional effects on multiple genes, our results cannot be unambiguously attributed to a specific function of Ube3a. Mechanistic inferences are further compounded by uncertainty regarding the ligase activity of the C-terminal flagged Ube3a copies overexpressed in our mouse model (Smith et al., 2011): whilst *in vitro* studies showed loss of ligase function upon C-terminal Ube3a tagging (Salvat et al., 2004; Kühnle et al., 2013; Avagliano Trezza et al., 2021; Bossuyt et al., 2021; Weston et al., 2021), *in vivo* investigations revealed largely increased ubiquitination levels in brain lysates from Ube3a2X mice (Khatri et al., 2018) and comparable behavioral phenotypes when C- or N-terminal tagged Ube3a is overexpressed in this model (Krishnan et al., 2017). Because Angelman syndrome (AS) reflects impaired Ube3a ligase function (Cooper et al., 2004), our finding that the transcriptional profile of Ube3a2X mice exhibits robust overlap with 15qdup, but not with AS, argues against a prominent loss of ligase activity in our model, and suggests that that the reported phenotypes may primarily reflect Ube3a-mediated transcriptional dysregulation. This hypothesis is consistent with Ube3a’s transcriptional effects being independent of its ligase activity (Nawaz et al., 1999; Li et al., 2006) and our observations of robust multiomic sex biasing influences, an effect that could reflect transcriptional coactivation with steroid hormone receptors (El Hokayem and Nawaz, 2014). Our finding that cortical transcriptome in Ube3a2X mice overlaps with gene lists related to steroid hormone receptor-relevant mechanisms lends further indirect support to this notion. Collectively, these observations suggest that *Ube3a* over-expression may impact gene networks and systems under the influence of diverse sex-relevant mechanisms, including the X chromosome, effects of genes sensitive to steroid hormone influence, downstream targets of steroid hormone receptors, as well as genes that are sex-differentially targeted by transcription factors. However, a putative downstream involvement of the ligase activity of Ube3a may also contribute to the sex-specific phenotypes we observed, since Ube3a has also been shown to ubiquitinate ER-α to target the receptor to proteasomal degradation (Gao et al., 2005). Future investigations of sex bias in other rodent models of *Ube3a* overdosage (Nakatani et al., 2009; Copping et al., 2017; Punt et al., 2022) may help corroborate or disprove these postulated mechanism.

The sex-specific effects observed in this study are congruent with both female-protective and male-enhancing risk explanations (Lai et al., 2015; Werling et al., 2016). *Ube3a* overexpression causes sex-specific transcriptional effects in many autism-associated genes, including *FMR1, SCN2A, PTEN, MEF2C, SOX5, GABRB2*, and *GABRB3* (Bourgeron, 2015). Many of these genes tend to be associated with autism via rare loss-of-function *de novo* mutations. Congruent with an interpretation of male-biased risk and female protection, these and other genes were underexpressed in Ube3a2X males, and over-expressed in females. Behaviorally, the observation that male Ube3a2X mice showed increased stereotyped behaviors, while no differences were apparent in females, is broadly consistent with a possible female protection, and male-potentiated genetic risk. It should however be noted that connectivity alterations in hypothalamic and motor-sensory areas were observed in female Ube3a2X mice. The possibility that this endophenotype is not compensatory, but instead the expression of a distinct etiopathological signature cannot be entirely ruled out. This notion would be consistent with emerging evidence that some of the autism-associated genes that are dysregulated in our model (e.g. *Shank3*, *Tsc2, Mef2c*, or *Fmr1*)can lead to pathological cascades of translational relevance when they are either under- or overexpressed during development (Oostra and Willemsen, 2003; Auerbach et al., 2011; Han et al., 2013).

Our findings also indicate that *Ube3a* overdosage results in sex-specific dysregulation of processes and pathways on which diverse autism-associated genetic influences have been theorized to converge (e.g., synaptic dysregulation, aberrant transcription and translation/protein synthesis, altered PIK3-AKT-mTOR signaling) (Bourgeron, 2015; Courchesne et al., 2019; Eyring and Geschwind, 2021). Diverse cell types theorized to be important in autism (e.g., excitatory and inhibitory neurons, glia cells) and which are affected by sex-relevant mechanisms are also impacted differentially by DE+ and DE-gene sets (Hu et al., 2011; Sarachana and Hu, 2013; Quartier et al., 2018; Velmeshev et al., 2019; Lombardo et al., 2020; Satterstrom et al., 2020; Trakoshis et al., 2020; Willsey et al., 2021; Gegenhuber et al., 2022; Kelava et al., 2022). Thus, over and above providing a sex-specific influence on key autism-associated genes, *Ube3a* overdosage may be changing these emergent processes/pathways and cell types in males versus females to confer heightened male-risk and female protection.

In conclusion, our data reveal robust sex-biasing effects on connectomics, repetitive behavior and transcriptomic organization in mice with extra copies of *Ube3a.* These results suggest that *Ube3a* can critically contribute to sex-bias in neurodevelopmental conditions like autism via influence on sex-relevant mechanisms, diverse neuronal and glial cell types, and important final common pathways that alter, synaptic organization, transcription, translation, and other key signaling pathways (e.g., PIK3-AKT-mTOR).

## Materials and Methods

### Ethical statement

Animal studies were conducted in accordance with the Italian Law (DL 26/2014, EU 63/2010, Ministero della Sanità, Roma) and the recommendations in the *Guide for the Care and Use of Laboratory Animals* of the National Institute of Health. Animal research protocols were also reviewed and approved by the Animal Care Committee of the University of Trento, Istituto Italiano di Tecnologia and the Italian Ministry of Health (authorization no. 560/16). All surgical procedures were performed under anesthesia.

### Animals

Mice were housed under controlled temperature (21 ± 1 °C) and humidity (60 ± 10%). Food and water were provided *ad libitum.* Generation of Ube3a2X mice (FVB/NJ background) was previously described in (Smith et al., 2011). Mice were purchased from Jackson Laboratory, Stock No: 019730. All experiments were performed on adult mice (< 12 months old) expressing two copies of *Ube3a* transgene (Ube3a2x), verified by quantitative PCR. Wild-type (WT) adult littermate mice were used as control.

### Resting state fMRI (rsfMRI)

rsfMRI was performed on four cohorts of mice: WT control females (n = 10); WT control males (n = 10); Ube3a2X females (n = 10); Ube3a2X males (n = 10). rsfMRI data were acquired as previously described (Ferrari et al., 2012; Sforazzini et al., 2016; Bertero et al., 2018; Pagani et al., 2019). Briefly, animals were anaesthetized with isoflurane (5% induction), intubated and artificially ventilated (2% maintenance during surgery). The left femoral artery was cannulated for continuous blood pressure monitoring. After surgery, isoflurane was replaced with halothane (0.7%) to obtain light sedation. Functional data acquisition started 45 min after isoflurane cessation.

Data were acquired with a 7T MRI scanner (Bruker) as previously described (Liska et al., 2015; Montani et al., 2020), using a 72-mm birdcage transmit coil and a 4-channel solenoid coil for signal reception. Co-centered single-shot rsfMRI time series were acquired using an echo planar imaging (EPI) sequence with the following parameters: TR/TE = 1000/15 ms, flip angle 30°, matrix 100 × 100, field of view 2.3 × 2.3 cm, 18 coronal slices, slice thickness 600 μm for 1920 volumes (total duration 32 minutes).

Mean arterial blood pressure (MABP) was recorded throughout the imaging sessions (Figure S6a-c). Ube3a2X had slightly lower MABP than control mice (2-way ANOVA, genotype effect, p < 0.05, Figure S6c), but values were well within the autoregulation window within which changes in peripheral blood pressure do not result in fMRI BOLD changes (Gozzi et al., 2007). In keeping with a negligible contribution of genotype-dependent MABP changes to our findings, we did not find any correlation between fMRI global connectivity and MABP in areas exhibiting sexspecific differences such as the PFC (r = −0.07,p = 0.67; Figure S6d). Arterial blood gas levels (pCO2 and pO2) were measured at the end of the acquisitions to ensure effectiveness of artificial ventilation. All mice had values within physiological range (pCO2 <42, pO2 > 90 mmHg, data not shown).

Analysis of body weight revealed, an effect of sex (p<0.01, 2-way ANOVA), but no genotype or sex*genotype interactions, p > 0.37, all tests; Figure S6e).

### rsfMRI connectivity mapping

Raw rsfMRI data were preprocessed as previously described (Sforazzini et al., 2014; Pagani et al., 2019; Montani et al., 2020). The initial 50 volumes of the time series were removed to allow for signal equilibration. Data were then despiked, motion corrected and spatially registered to a common reference mouse brain template. Motion traces of head realignment parameters (3 translations + 3 rotations) and mean ventricular signal (corresponding to the averaged BOLD signal within a reference ventricular mask) were regressed out from each time course. All rsfMRI time series were also spatially smoothed (full width at half maximum of 0.6 mm) and band-pass filtered to a frequency window of 0.01-0.1 Hz.

To obtain an unbiased identification of the brain regions exhibiting alterations in functional connectivity, we calculated global local fMRI connectivity maps for all mice. Global fMRI connectivity is a graph-based metric that defines connectivity as the mean temporal correlation between a given voxel and all other voxels within the brain. Local connectivity strength was mapped by limiting this measurement to connections within a 0.6252 mm (6 voxels in-plane) sphere around each voxel (Cole et al., 2010; Liska et al., 2015). rsfMRI connectivity was also probed using a seed-based approach (Montani et al., 2020; Rocchi et al., 2022). A 3×3×1 seed region was selected to cover the areas of interest and VOI-to-seeds correlations were computed. Pearson’s correlation scores were first transformed to z-scores using Fisher’s r-to-z transform and then averaged to yield the final connectivity scores.

Voxel-wise intergroup differences in global and local connectivity and seed-based maps were assessed using a linear model including sex, genotype and sex*genotype as factors (lm function in R studio). Data was imported into R using the oro.nifti package. The obtained t score maps were (FWER) cluster-corrected using a cluster threshold of p = 0.05.

### Structural MRI

To locate and quantify gray matter (GM) changes in Ube3a2X mice, we performed postmortem voxel-based morphometry (VBM) as previously described (Pagani et al., 2016b). Briefly, mice were deeply anesthetized with 5% isoflurane, and their brains were perfused via cardiac perfusion of 4% PFA added with a gadolinium chelate to shorten longitudinal relaxation times. High-resolution morpho-anatomical T2-weighted MR imaging of mouse brains was performed using a 72 mm birdcage transmit coil, a custom-built saddle-shaped solenoid coil for signal reception. For each session, high-resolution morpho-anatomical images were acquired with the following imaging parameters: FLASH 3D sequence with TR =17 ms, TE =10 ms, α= 30°, matrix size of 260 x 180 x 180, FOV of 1.83 x 1.26 x 1.26 cm, and voxel size of 70 um (isotropic).

Morpho-anatomical differences in local GM volumes were mapped using a registration-based VBM procedure (Pagani et al., 2016b, 2019; Pucilowska et al., 2018). Specifically, high-resolution T2-weighted images were corrected for intensity nonuniformity, skull stripped, and spatially normalized to a study-based template using affine and diffeomorphic registrations. Registered images were segmented to calculate tissue probability maps. The separation of the different tissues is improved by initializing the process with the probability maps of the study-based template previously segmented. The Jacobian determinants of the deformation field were extracted and applied to modulate the GM probability maps calculated during the segmentation. This procedure allowed the analysis of GM probability maps in terms of local volumetric variation instead of tissue density. Brains were also normalized by the total intracranial volume to further eliminate overall brain volume variations and smoothed using a Gaussian kernel of 3 voxel width. To quantify volumetric changes identified with VBM, we used preprocessed images to independently calculate the size of neuroanatomical areas via volumetric anatomical labelling (Pagani et al., 2016b).

### Behavioral tests

#### Open field test

To test spontaneous locomotion, experimental mice were individually placed in an open field arena (40 cm × 40 cm × 40 cm) and let free to explore for 10 minutes. The walls of the arena were smooth and gray colored. Sessions were recorded and mice were automatically tracked using EthoVisionXT (Noldus). Locomotor activity was measured as total distance and mean velocity. In addition, the proportion of time spent in the center of the arena and outer zones was analyzed to estimate the level of anxiety. Number of full body rotations, time spent wall rearing and selfgrooming were used as measures of stereotypic behavior.

#### Rotarod

The rotarod test is widely used for the evaluation and assessment of locomotor activity and motor coordination in rodents (Crawley, 1999). Mice were pre-trained on the rotarod apparatus for 3 days before the test. This habituation step consisted in a performance at a constant speed of 4 rpm for 5 minutes. On the fourth day, mice were tested for 5 minutes in 3 different trials. During each trial, the rotating rod accelerated from 4 to 64 rpm. Mice had 5 minutes of rest between each trial. The total time that the mice spent on the rotating rod was measured. The trials ended when the mice fell down or 3 consecutive full rotations were observed.

#### Habituation/dishabituation social interaction test

Animals were tested as previously described (Huang et al., 2014). Experimental mice were individually placed in a testing cage (GR900 Tecniplast cages (904 cm2)), lightly illuminated (5±1 lux)), 1 hour before the test. A matching stimulus mouse (same sex, same strain and same age) was introduced into the testing cage for a 1-minute interaction. At the end of the trial, the stimulus mouse was removed for 3 minutes. This sequence was repeated for 4 trials. Finally, experimental mouse was tested in a fifth 1 min-dishabituation trial where a new stimulus mouse was introduced in the testing cage. Time spent interacting (sum of nose-to nose sniffing, anogenital sniffing and following) was scored across trials by an experimenter blind to genotypes.

#### Three-Chamber Social Interaction Test

Each mouse’s preference for a conspecific over an inanimate object (sociability), as well as its preference for a stranger mouse over a familiar mouse (social novelty) was assessed using previously established 3-chamber assays (Silverman et al., 2010b). During the sociability phase, a stranger mouse was placed in one chamber inside a wire cup that allowed nose contact. Video monitoring of the test mouse’s exploration of the apparatus was carried on for 10 min. Next, in the social novelty phase, the test mouse was re-exposed for 10 min to the initial stranger (now familiar) mouse, as well as to a novel stranger mouse placed inside the second wire cup, in the opposite chamber. The amount of time spent sniffing the wire cups was measured. A sociability index was calculated as the time spent with the mouse cup minus time spent with the empty cup divided by the total time of investigation 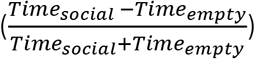
. Accordingly, the sociability index during the social novelty phase (Novelty index) was calculated as the time spent with the novel mouse cup minus time spent with the familiar mouse cup divided by the total time of investigation 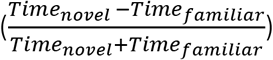
.

#### RNA-sequencing and preprocessing

Mice of both genotypes and sexes were sacrificed by cervical dislocation, and prefrontal cortex and hypothalamus were rapidly identified according to the Allen Mouse Brain Atlas (http://www.brain-map.org) and dissected. The samples used for RNA-sequencing (RNA-Seq) experiment are deposited in Gene Expression Omnibus (GEO) (GSE217420). For a complete list of quality control parameters for RNA-seq, please see Supplementary Table S4. Brains were rotated and the exposed hypothalamus was excised with surgical tweezers. To collect prefrontal cortex tissue, explanted brains were placed in an adult mouse brain matrix (Agnthos, Sweden). Two coronal sections at the level of the prefrontal cortex for each mouse brain were collected (Spijker, 2011). The sections (1mm thick) were cut with scalpel blades and immediately put on a semi-frozen glass slide. Tissue from PFC was obtained by micro punches of 0.5 mm. One micro punch for each hemisphere/section was collected, for a total of four micro punches per brain. Following tissue collection, samples were frozen with dry ice and stored at −80 °C until RNA extraction.

RNA Extraction and Library Preparation. All procedures were conducted in RNAase-free conditions. On the day of RNA extraction, the hypothalamus and PFC tissues were disrupted and homogenized for 3 minutes using motor-driven grinders. Total RNA isolation was performed using RNeasy Mini Kit and RNeasy Micro Kit (Qiagen) respectively, following manufacturer’s instructions. RNA concentration was evaluated using Qubit RNA BR Assay Kit (Life Technologies). RNA purity was assessed by determining UV 260/280 and 260/230 absorbance ratios using a Nanodrop^®^ ND-1000 spectrophotometer (Thermo Fisher Scientific). RNA quality was evaluated by measuring the RNA integrity number (RIN) using an Agilent RNA 6000 Nano Kit with an Agilent 2100 Bioanalyzer (Agilent Technologies, Santa Clara, CA, USA) according to the manufacturer’s instructions. All samples had RIN of > 6.8 (see Table S4). Libraries for RNA-seq were prepared using the paired-end TruSeq^®^ Stranded mRNA Sample Preparation kit (Illumina, San Diego, Ca, USA) according to manufacturer’s instructions. For each sample of hypothalamus and PFC, 1000 ng and 500 ng were used as input quantity, respectively. The libraries were prepared in one batch using NovaSeq 6000 S2 Reagent Kit (200 cycles) at an average read-depth of 100 million paired-end reads. Libraries were 0.85 nM in a volume of 150 μl and loaded on an Illumina NovaSeq 6000 System (IIT-Center for Human Technologies - Genomic Unit-Genova GE - IT).

Raw reads were aligned to the mm10 genome (GRCm38 primary assembly obtained from the Gencode website) usign the STAR aligner and counted with featureCounts using the gene annotation Gencode v24. Picard tools (http://broadinstitute.github.io/picard/) functions were used to quantify sequencing-related variables. Low read genes were removed if they had less than 100 reads in 2 or more samples. Variance filtering was used to filter out genes in the bottom 15%-tile ranked by variance. In PFC samples, this filtering resulted in 12,294 genes being retained for further analysis, while 13,727 genes were retained for Hypo samples. Normalization for library size was implemented with the *calcNormFactors* function in the edgeR R library, using the trimmed mean of M values (TMM) method (Robinson and Oshlack, 2010). The *voom* function from the *limma* library in R (Law et al., 2014) was then used to transform the data to log counts per million and estimate precision weights to incorporate in the linear modeling of differential expression (DE). Finally, surrogate variable analysis (SVA) was used to estimate artifact-related variables (Leek et al., 2012). This was achieved by constructing a model of the known variables to account for (i.e. sex, genotype, sex*genotype interaction, and RIN), and then having SVA estimate surrogate variables (SV) from the error term of the model. The number of surrogate variables (SV) estimated was 2 for both PFC and Hypo. In order to better understand how these SVs account for known sequencing-related artifacts variables we first ran a PCA on the Picard variables of percent coding bases, percent utr bases, percent intronic bases, percent intergenic bases, median CV coverage, median 5’ to 3’ bias, aligned reads, and AT dropout. This gave an orthogonal summary of the artifact-related variables, and we then analyzed these PCs for correlations with the SVs. We find that both SVs are highly correlated with many of the sequencing-related artifact PCs (Supplementary Figure S7), indicating that substantial sequencing-related artifact is accounted for parsimoniously with 2 SVs. Furthermore, to see how these SVs correlate with principal axes of variance in the expression data, we correlated SVs with the first 10 principal components of the expression data. Here again, the SVs known to be relevant for sequencing-related artifact are highly correlated with the first PCs in the expression data (Supplementary Figure S7), indicating that without removing these significant artifact-related structured noise variables, they would swamp a large proportion of the variance in the expression data. Therefore, we used these SVs in the linear modeling for differential expression to account for and remove variance associated with these artifact-related SVs.

#### Differential expression analysis

Differential expression (DE) analysis was achieved using functions for linear modeling in the *limma* library in R. DE analysis examined specific contrasts for the sex*genotype interaction as well as main effects of sex and group respectively. The linear model included RIN and SVs as covariates and incorporated the precision weights estimated by *voom.* DE models were computed separated for PFC and Hypo samples. Genes that pass FDR q<0.05 (Storey, 2002) were considered DE.

#### Enrichment analysis

To annotate DE gene sets for KEGG pathways and mouse brain cell types, we used the *enrichR* library in R (https://maayanlab.cloud/Enrichr/) (Chen et al., 2013; Kuleshov et al., 2016). Mouse brain cell types were based on data cortex and hippocampal tissue samples in the Allen Institute 10x single-cell RNA-seq dataset (Yao et al., 2021) (https://celltypes.brain-map.org/rnaseq/mousectx-hip10x). For Gene Ontology biological process (GO BP) enrichment analysis, we utilized the *GeneWalk* library in Python (https://github.com/churchmanlab/genewalk) (Ietswaart et al., 2021). With custom gene lists of relevance to autism- and sex-related genomic mechanisms, we ran additional gene set enrichment tests with DE gene sets. These tests were run using custom code for running gene set enrichment analysis that computes enrichment odds ratios and p-values based on the hypergeometric distribution. For these tests, we used a background total equivalent to the total number of genes analyzed after filtering in the main gene expression analyses (e.g., 12,294 for PFC and 13,727 for Hypo). Because autism- and sex-related gene lists are based on human gene symbols, we first converted mouse gene IDs into human gene homologs and then ran all enrichment tests. All results of these enrichment tests were thresholded at FDR q<0.05. The diagnostic gene lists were a curated list of high-impact autism-associated genes from SFARI gene (https://gene.sfari.org; downloaded Jan, 2021) and differentially expressed genes from post-mortem cortical tissue in autism, schizophrenia, bipolar disorder (Gandal et al., 2018), and duplication 15q syndrome (Parikshak et al., 2016) and from iPSC-derived neurons from dup15q and Angelman syndrome patients (Germain et al., 2014). Sex-related gene lists included downstream targets of the androgen (Quartier et al., 2018) and estrogen receptors (Gegenhuber et al., 2022), and genes that are sex differentially targeted by transcription factors (Lopes-Ramos et al., 2020). We also ran similar enrichment analyses for chromosomes in order to test if DE gene sets were significantly enriched for genes located on specific chromosomes (e.g., X chromosome).

#### Protein-protein interaction analysis

To better understand how key DE genes may work together in specific ASD-relevant systems biological processes, we used STRING-DB (https://string-db.org/) to conduct a proteinprotein interaction (PPI) analysis whereby the input gene list were *UBE3A*, steroid hormone receptors (AR, ESR1, ESR2), and PFC sex-by-genotype DE genes that were annotated as significantly enriched in GeneWalk GO biological processes of relevance to autism (e.g., synaptic, transcription, translation, mTOR, and ERK signaling pathways) or of relevance to steroid hormone receptor signaling. This analysis was done using the human gene homologs of the mouse DE genes, and applies standard STRING defaults in the analysis (i.e. full network type, confidence level = 0.4). The resulting PPI network plot is then colored with a data-driven k-means clustering with k=3 in order to visually demarcate proteins that cluster into largely synaptic, transcription/signaling, and translation sets.

## Code availability

The code used for preprocessing and analyzing mouse rsfMRI data is available at https://github.com/functional-neuroimaging/rsfMRI-preprocessing https://github.com/functional-neuroimaging/rsfMRI-global-local-connectivity https://github.com/functional-neuroimaging/rsfMRI-seed-based-mapping Code for RNA-seq analyses is available at https://github.com/IIT-LAND/ube3a_rnaseq

## Acknowledgments

This work was supported by Simons Foundation Grants (SFARI 400101) to A. Gozzi and the European Research Council (ERC—DISCONN, No. 802371). A. Gozzi was also supported by Brain and Behavior Foundation 2017 (NARSAD—National Alliance for Research on Schizophrenia and Depression), NIH (1R21MH116473-01A1), the Telethon foundation (GGP19177). M. Pagani was supported by the European Union’s Horizon 2020 research and innovation programme under grant agreement No. 845065 (Marie Sklodowska-Curie Global Fellowship - CANSAS). M.V. Lombardo acknowledges funding by the European Research Council (ERC) under the European Union’s Horizon 2020 research and innovation programme under grant agreement No 755816. GP was supported by the Brain and Behavior Research Foundation (NARSAD Young Investigator Grant; ID: 26617) and the University of Trento (Starting Grant for Young Researchers). Y. B. was supported by TRAIN (Trentino Autism Initiative), a strategic project of the University of Trento.

## Notes

### Competing Interest Statement

The authors have declared no competing interest.

### Summary of Updates

This revision incorporates new transcriptional enrichment data and extended discussion

