## Supplementary Figures for "Sex-biasing influence of autism-associated *Ube3a* gene overdosage at connectomic, behavioral and transcriptomic levels"

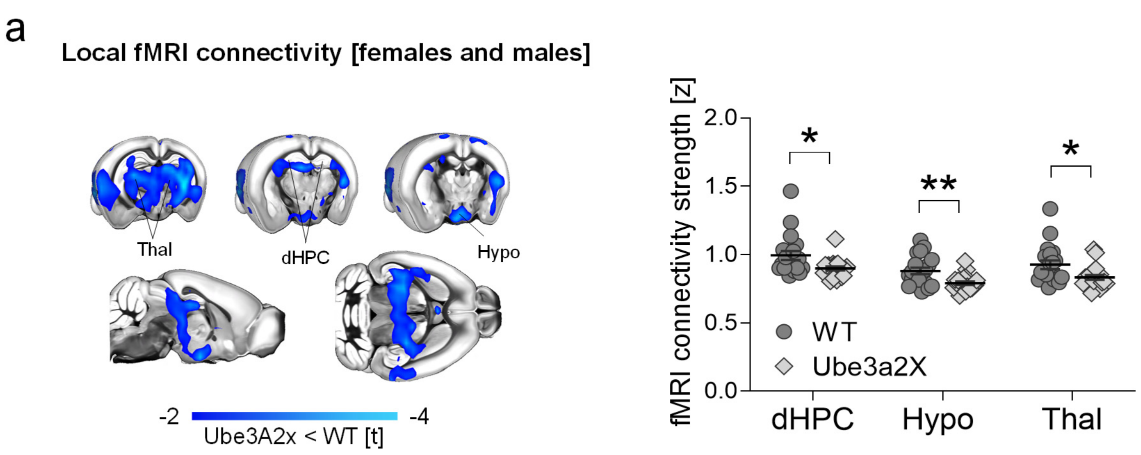


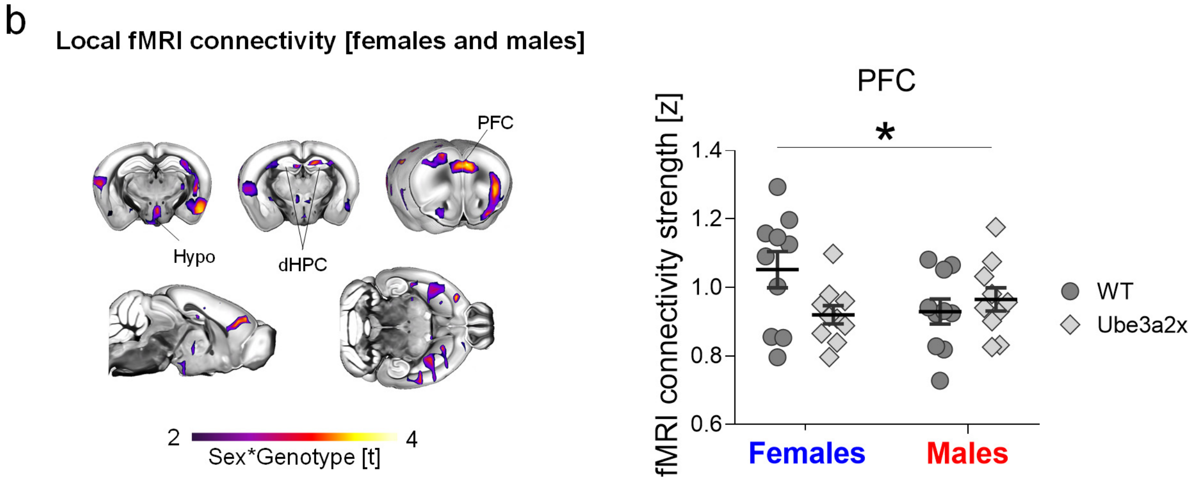


Supplementary Figure S1

Increased *Ube3a* dosage affects local fMRI connectivity in a sex-dependent manner. a) Intergroup contrast maps (left panel) showing reduced local fMRI connectivity strength in Ube3a2X animals (n = 20) compared to WT control (n = 20) littermates (both sexes, blue color indicates reduced connectivity, t-test, t > 2; FWE cluster-corrected). Panel on the right illustrates quantification of global fMRI connectivity strength in representative regions of interest (t-test, thalamus, t = 2.57, p = 0.015; dorsal hippocampus, t = 2.62, p = 0.014; hypothalamus, t = 3.25, p = 0.028). b) Contrast maps (left panel) showing areas exhibiting sex*genotype interactions in local fMRI connectivity strength (purple and yellow coloring, t > 2; FWE cluster-corrected). Panel on the right illustrates quantification sex*genotype interaction in representative regions of interest. (ANOVA interaction, F = 6.68, p = 0.019). dHPC, dorsal hippocampus; Hypo, Hypothalamus; PFC, Prefrontal Cortex, Thal, Thalamus. *p < 0.05, **p < 0.01. FWE: family-wise error. Error bars of the plots indicate SEM and each dot represents a mouse.

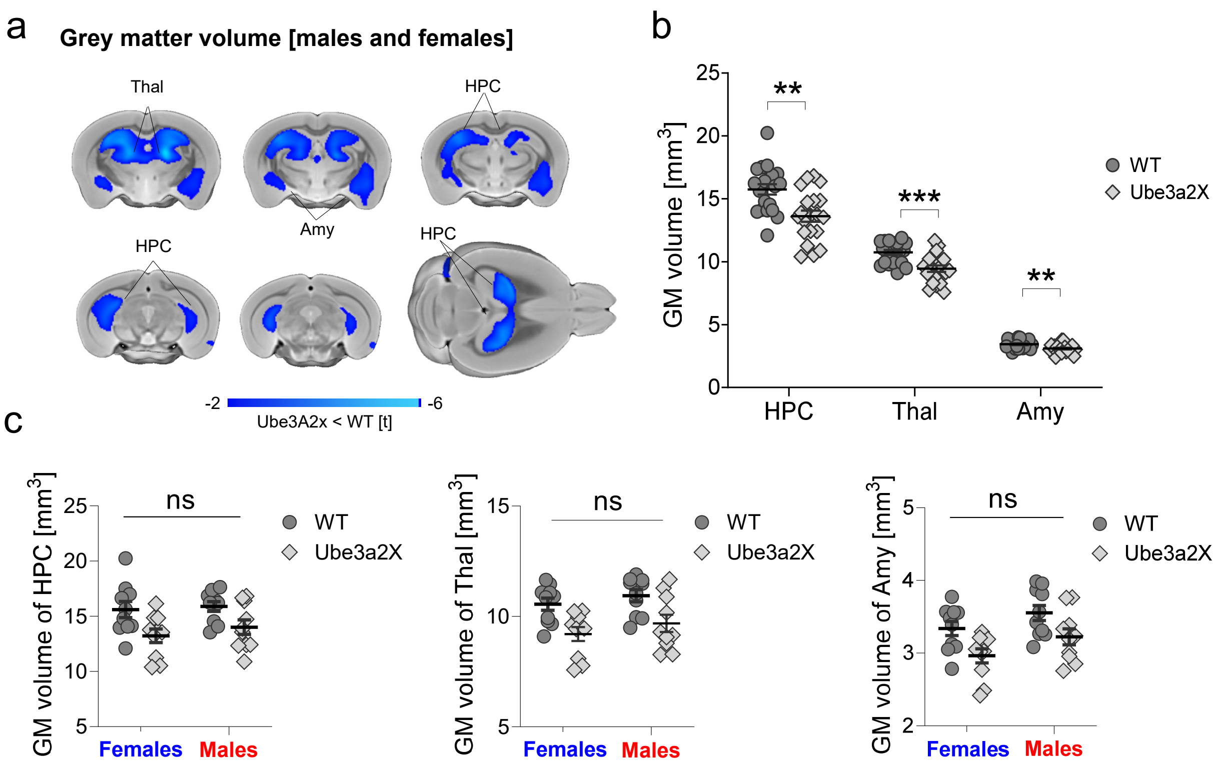


Supplementary Figure S2

**Brain anatomy is not affected by *Ube3a* dosage in a sex-dependent manner. a)** Structural MRI showing a reduction in gray matter volume in Ube3a2X mutants (n=20) compared to WT control (n=20) littermates (both sexes, t > 2; FWE cluster-corrected, blue coloring). **b)** Regional quantifications showing reduction of gray matter volume in hippocampus (t = 3.47, p = 0.001), thalamus (t = 4.097, p < 0.001) and amygdala (t = 3.29, p = 0.002) in Ube3a2X mutants compared to WT mice (both sexes). **c)** Sex*genotype interaction in gray matter volume was not significant ("ns") in all the quantified regions, including amygdala (ANOVA's interaction, F = 0.05, p = 0.82), thalamus (ANOVA's interaction, F = 0.025, p = 0.87) and hippocampus (ANOVA's interaction, F = 0.17, p = 0.68). The absence of significant sex*genotype interactions reflects concomitantly reduced GM volume in both Ube3a2X females (t-test, amygdala, t = 2.59, p = 0.027; thalamus, t = 3.60, p = 0.009; hippocampus, t = 2.71, p = 0.002) and Ube3a2X males (unpaired t-test, amygdala, t = 2.59, p = 0.057; thalamus, t = 2.78, p = 0.017; hippocampus, t = 2.13 , p = 0.08) compared to sex-matched WT control mice. Amy, Amygdala; Thal, Thalamus; Hypo, Hypothalamus; *p < 0.05, **p < 0.01; FWE, family-wise error; ns, non-significant; GM, gray matter, Error bars of the plots indicate SEM and each dot represents a mouse.

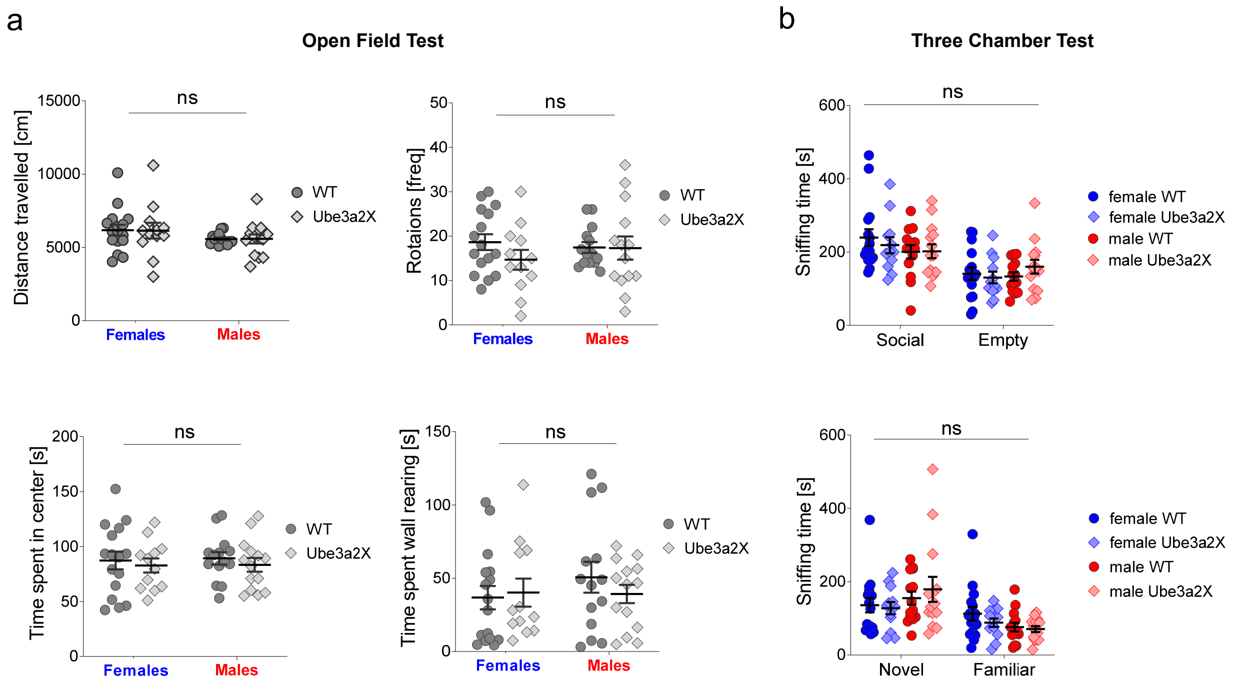


Supplementary Figure S3

Ube3a2X mice do not show gross alterations in social and anxiety-related behaviors. a) Results of open field test. Sex*genotype interaction was not significant (“ns”) for travelled distance ($F = 0.005$, $p = 0.94$), frequency of rotations ($F = 0.89$, $p = 0.35$), total time spent in the center ($F = 0.011$, $p = 0.92$) and wall rearing ($F = 0.73$, $p = 0.39$, ANOVA’s interaction). All four behaviors were unimpaired in Ube3a2X mutants (n = 27, n = 14 males and n = 13 females) compared to sex-matched WT littermates (n = 30, n = 14 males and n = 16 females) ($p > 0.20$, all behaviors). b) Three chamber test. The total time spent sniffing is reported for sociability (top panel) and the social novelty phase (bottom panel). Sex*genotype interaction in sociability (ANOVA’S interaction, $F = 1.24$, $p = 0.31$) and preference for social novelty (ANOVA’s interaction, $F = 0.39$, $p = 0.76$) were not significant (“ns”). Chamber effect was significant in both phases ( $F = 20.71$, $p < 0.0001$, $F = 18.3$, $p < 0.0001$, respectively). ***$p < 0.001$. All plots report mean± SEM, on top of individual data points.

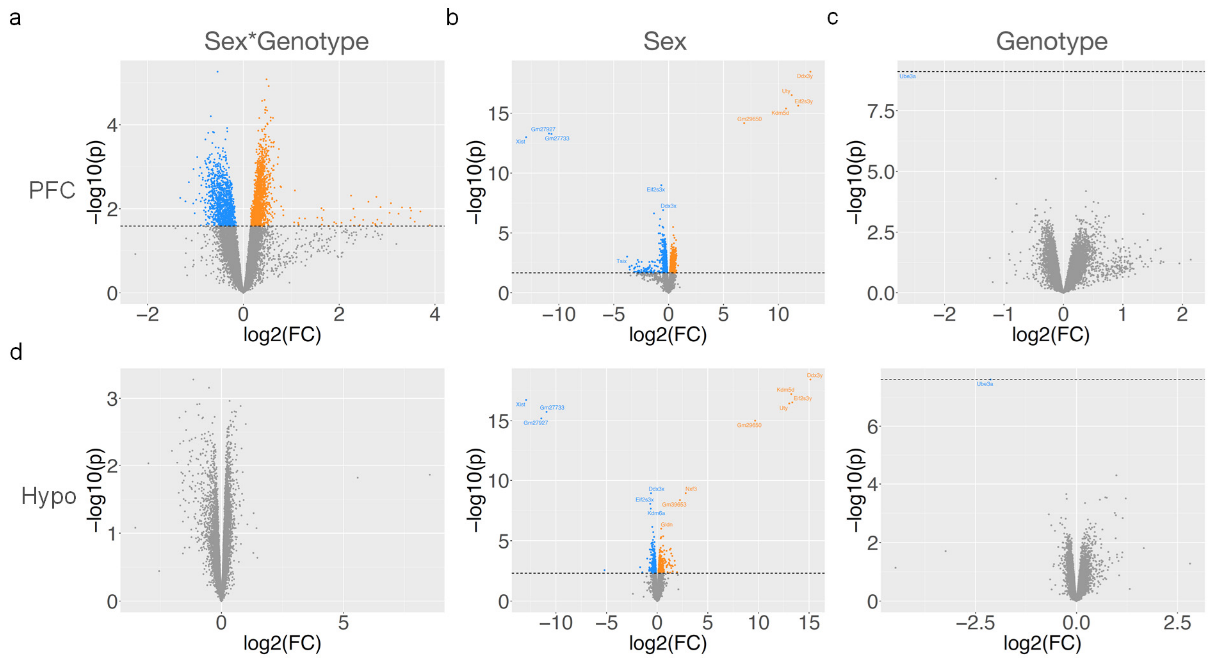


Supplementary Figure S4

**Genes are dysregulated in a sex*genotype manner in the PFC.** Volcano plots show log2 fold change (FC) on the x-axis and -log10 p-values on the y-axis for the effects of the sex*genotype interaction (left), main effect of sex (middle), and main effect of genotype (right). Rows indicate PFC data (top) or Hypo data (bottom). Genes colored in orange are the DE- genes, while genes colored in blue are DE+ genes. Genes depicted in gray fall below the horizontal dotted line indicating the FDR q<0.05 threshold, and are thus considered not differentially expressed.

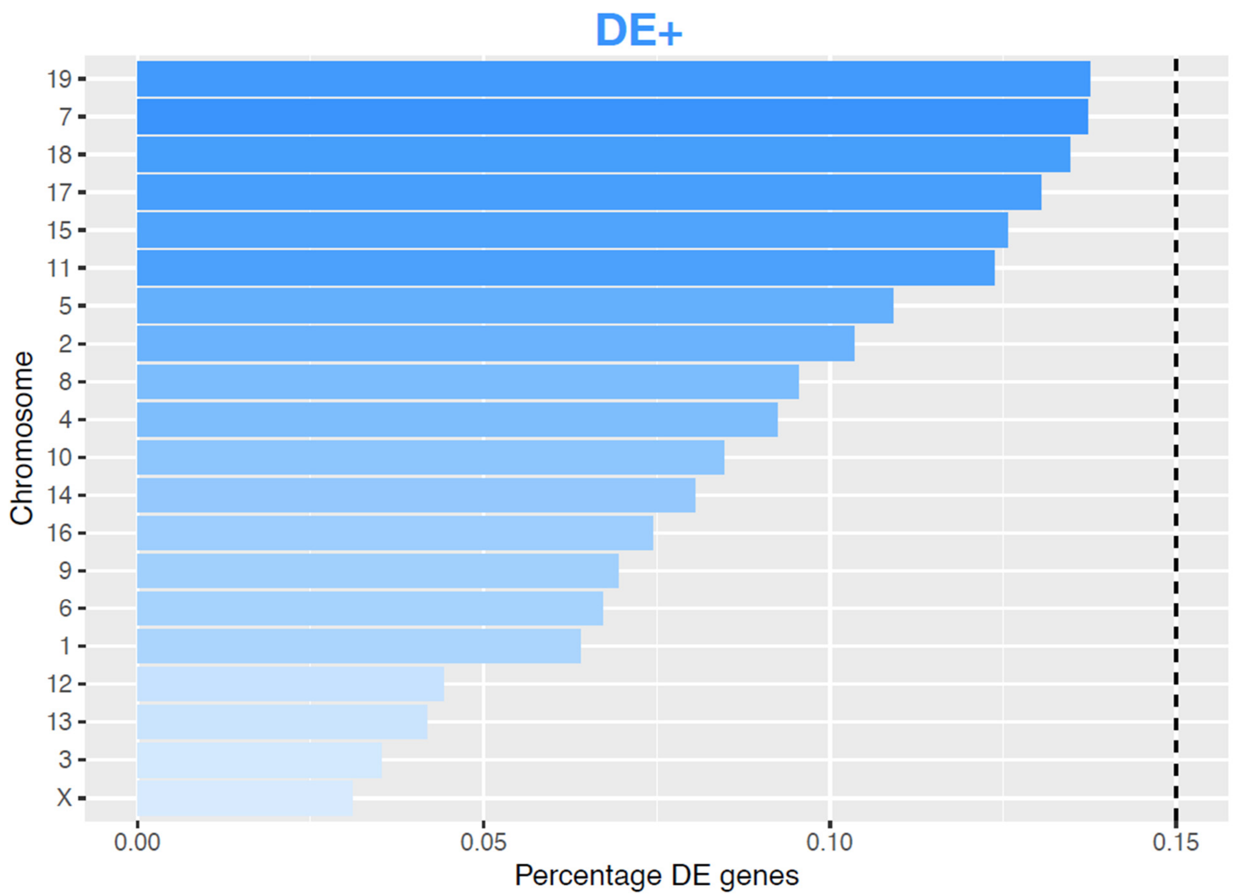


Supplementary Figure S5

**DE+ genes are equally distributed across chromosomes.** Plot showing the percentage of genes per each chromosome that are DE+. Color indicates the enrichment odds ratio. The vertical dotted line indicates the FDR q < 0.05 threshold.

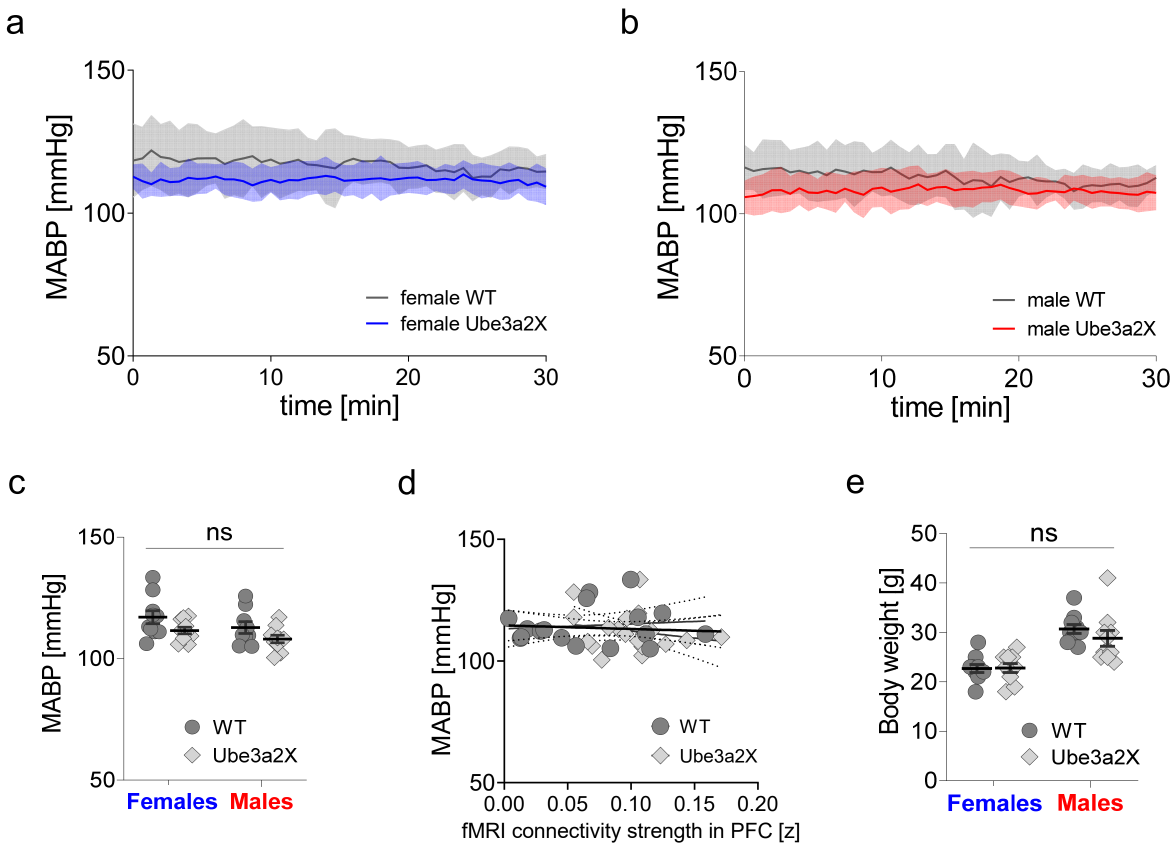


Supplementary Figure S6

**Mean arterial blood pressure and body weight.** a) Plot of mean arterial blood pressure in female Ube3a2X mice (n=10) and female WT littermates (n=10) during rsfMRI scanning. b) Plot of arterial blood pressure in male Ube3a2X mice (n=10) and male WT littermates (n=10) during rsfMRI scanning. c) Quantification of mean arterial blood pressure of Ube3a2X females (unpaired t-test, t = 1.91, p = 0.12) and Ube3a2X males (unpaired t-test, t = 1.62 p = 0.21) across the imaging time window. No significant sex*genotype interaction was observed in arterial blood pressure (ANOVA's interaction, F = 0.031, p = 0.86, "ns" in the plot). d) Lack of correlation between fMRI connectivity in a representative brain region (i.e. the prefrontal cortex) and mean arterial blood pressure (r = -0.07, p = 0.67). e) Body weight of Ube3a2X females (unpaired t-test, t = 0.06, p = 0.99) and Ube3a2X males (unpaired t-test, t = 1.22 p = 0.40) is comparable to that of sex-matched WT control mice. Sex*genotype interaction in body weight was not statistically significant (ANOVA's interaction, F = 0.82, p = 0.37, "ns" in the plot). ns, non-significant. Error bars of the plots indicate SEM.

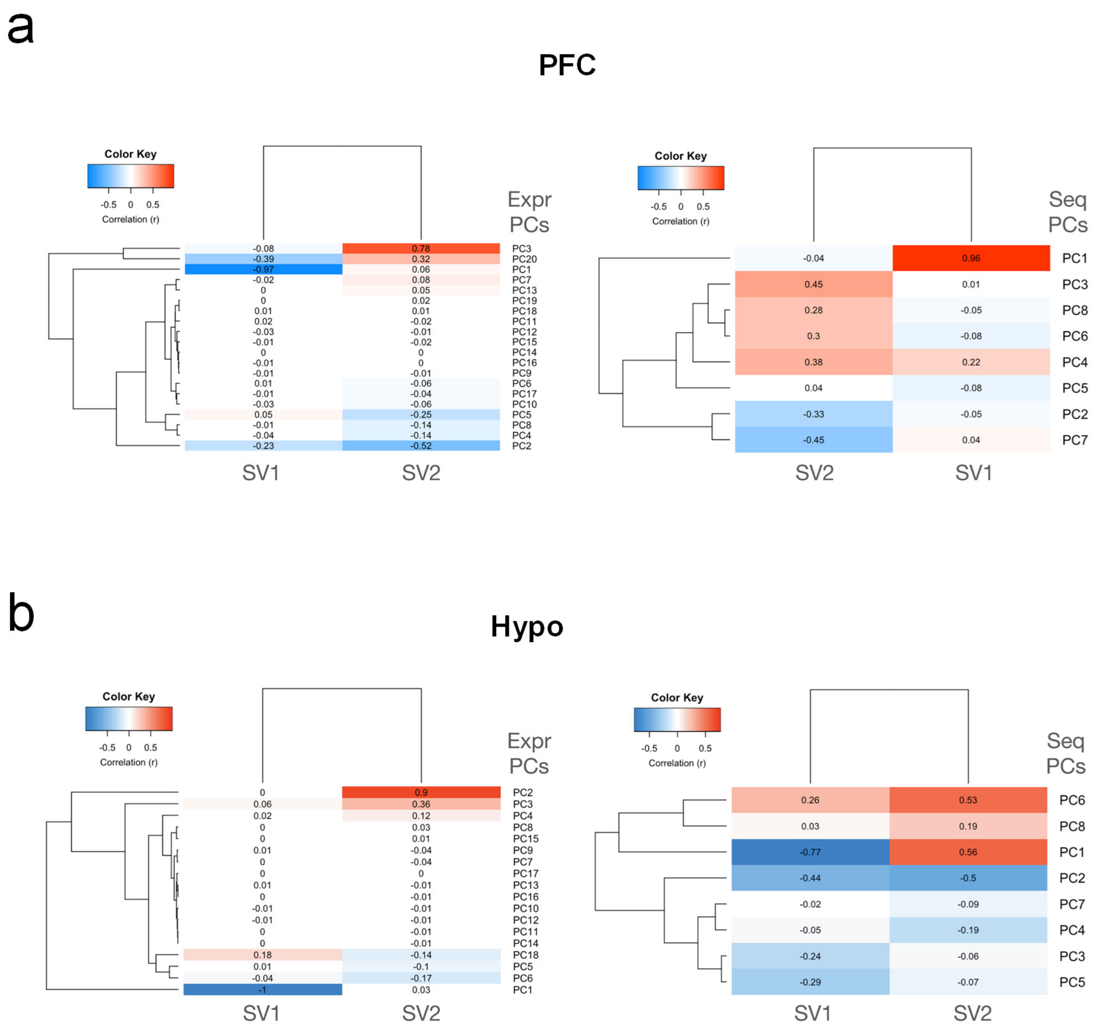


Supplementary Figure S7

Correlations between surrogate variables (columns SV1, SV2) and principal components (rows) derived from preprocessed gene expression data (left) or sequencing-related variables (right). Panel (a) shows these correlations for PFC data, while panel (b) shows the correlations from Hypo data.
